# Distinct components of nucleoside-modified messenger RNA vaccines cooperate to instruct efficient germinal center responses

**DOI:** 10.1101/2024.05.17.594726

**Authors:** Emily Bettini, Aleksey Chudnovskiy, Giulia Protti, Sandra Nakadakari-Higa, Simona Ceglia, Diana Castaño, Joy Chiu, Hiromi Muramatsu, Thandiswa Mdluli, Edit Abraham, Zoltan Lipinszki, Ivan Maillard, Ying K. Tam, Andrea Reboldi, Norbert Pardi, Roberto Spreafico, Gabriel D. Victora, Michela Locci

## Abstract

Nucleoside-modified mRNA vaccines elicit protective antibodies through their ability to promote T follicular helper (Tfh) cells. The lipid nanoparticle (LNP) component of mRNA vaccines possesses inherent adjuvant activity. However, to what extent the nucleoside-modified mRNA can be sensed and contribute to Tfh cell responses remains largely undefined. Herein, we deconvoluted the signals induced by LNP and mRNA that instruct dendritic cells (DCs) to promote Tfh cell differentiation. We demonstrated that the nucleoside-modified mRNA drives the production of type I interferons that act on DCs to induce their maturation and the induction of Th1-biased Tfh responses. Conversely, LNP favors the acquisition of a Tfh cell-inducing program in DCs, a stronger Th2 polarization in Tfh cells, and allows for rapid mRNA translation by DCs within the draining lymph node. Our work unravels distinct adjuvant features of mRNA and LNP necessary for the induction of Tfh cells, with implications for vaccine design.

## INTRODUCTION

The severe acute respiratory syndrome coronavirus 2 (SARS-CoV-2) pandemic propelled the approval of the first mRNA vaccines for clinical use. Since then, billions of doses of mRNA vaccines have been administered worldwide^1^, and this vaccine platform has proven to be highly successful at driving potent neutralizing antibody (nAb) and memory B cell (MBC) responses, as well as preventing severe coronavirus disease 2019 (COVID-19)^2–5^. Long-lived plasma cells (LLPCs), which secrete Abs, and MBCs are canonically generated through germinal center (GC) reactions^6^. It is within GCs that antigen-specific B cells undergo iterative rounds of somatic hypermutation, ultimately leading to the formation of affinity-matured MBCs and LLPCs. Previous work from our group and others has demonstrated that mRNA vaccines, comprised of nucleoside-modified and purified mRNA encapsulated in lipid nanoparticles (mRNA-LNP), drive potent GC responses in both mice and humans, which correlate with nAb production and antigen-specific MBC responses^7–10^. However, the mechanism by which mRNA-LNP instruct GC responses remains largely undefined.

GC reactions are regulated by a specialized subset of CD4 T cells referred to as T follicular helper (Tfh) cells^11–13^. Tfh cell differentiation is a multistep, multifactorial process that begins when antigen-presenting cells (APCs), often dendritic cells (DCs), present antigen to naïve CD4 T cells within the T cell zone of a secondary lymphoid organ. During this interaction, the naïve CD4 T cells can receive a multitude of pro-Tfh signals, including T cell receptor (TCR) stimulation, costimulatory molecules (CD80/CD86 and ICOS-L), and cytokines (IL-6 and IL-21 in mice, IL-12, activin A, and TGF-β in humans) from the APCs that prompt the initiation of Tfh cell differentiation^11,13,14^. Conversely, antagonistic signals driven by IL-2 and its high-affinity receptor CD25 can suppress the early differentiation of Tfh cells^15,16^. Activated CD4 T cells also upregulate the oxysterol receptor Ebi2, which allows for early positioning of these cells at the interface of the B cell follicles and T cell zone (T-B border), where they commit to the Tfh differentiation pathway by interacting with ICOSL^+^CD25^+^ DCs^17^. At T-B borders, important signals from cognate B cells also contribute to the subsequent differentiation into fully mature Tfh cells^13^.

We have previously demonstrated that the LNP component of mRNA vaccines has intrinsic adjuvant activity, which can favor Tfh cell differentiation through the induction of the pro-Tfh cytokine IL-6^8^. However, it is not completely understood whether signals important for the differentiation of Tfh cells also stem from sensing the nucleoside-modified mRNA. The introduction of modified nucleosides such as pseudouridine in the mRNA was a transformative finding originally introduced to restrain the capacity of Toll-like receptor (TLR) 3 and TLR7/8 to sense the *in vitro* transcribed mRNA and induce the expression of pro-inflammatory cytokines such as Tumor necrosis factor (TNF)-α and type I interferons (IFNs)^18–20^. The observation that the injection of nucleoside-modified mRNA *in vivo* does not prompt measurable type I IFN production in circulation^21^, suggests that this vaccine component is immunosilent. However, it has not been elucidated whether nucleoside-modified mRNA can trigger the local production of low levels of type I IFNs that are biologically relevant for Tfh cell responses. Our recent finding that mice lacking the TLR signaling adaptor MyD88 mount hindered Tfh cell responses following immunization with mRNA-LNP but not upon injection of protein subunit antigens mixed to empty LNP (protein-LNP)^8^, spurred the hypothesis that the N1-methylpseudouridine modification and stringent mRNA purification do not completely abolish the capacity of TLRs to recognize the mRNA component of mRNA-LNP vaccines.

Herein, we sought to uncouple the signals promoted by the LNP from those driven by the modified and purified mRNA that elicit optimal Tfh and GC responses. Our work uncovered the importance of type I IFNs produced in response to the mRNA component of mRNA-LNP vaccines in promoting DC maturation and enhancing the magnitude of GC reactions by acting directly on DCs. Of note, mRNA-driven type I IFN signaling in DCs drove Th-1 polarized CD4 T cell responses, while the LNP component promoted mixed Th1/Th2 responses more biased toward a Th2 profile. On the other hand, the sensing of LNP instructed a transcriptional program in DCs which included costimulatory molecules as well as chemoattractant receptors important for positioning at T-B borders, such as Ebi2. By using a LIPSTIC-based approach, we found a transcriptional signature associated with antigen presentation in DCs as well as diffuse DC activation following immunization with LNP-containing vaccines. This broad DC activation was explained by the finding that most conventional DCs in the draining lymph node (dLN) could efficiently internalize LNP and translate mRNA *in situ*, with only a small minority of DCs in the dLN that processed mRNA-LNP at the site of vaccine injection, as shown by photoactivation studies. Altogether, our work underscores a dual immunological mechanism of action of mRNA-LNP vaccines and unfolds the distinct adjuvant properties of the mRNA and LNP vaccine components.

## RESULTS

### Nucleoside-modified mRNA drives a type I IFN response that amplifies GC reactions

Unmodified phosphodiester RNA is recognized by TLRs and leads to the production of type I IFNs (IFN-α and IFN-β)^22–24^, which in turn, can enhance humoral immunity and Tfh cell responses by stimulating DC maturation and the capacity to produce the pro-Tfh cytokine IL-6^25–28^. Our unexpected finding that mice lacking the TLR adaptor MyD88 mounted dampened Tfh cell responses upon immunization with mRNA-LNP^8^ prompted us to hypothesize that, despite containing N1-methylpseudouridine, the mRNA component of mRNA-LNP vaccines can still be sensed by TLRs and have a biologically relevant impact on vaccine-driven B cell responses. Hence, we sought to determine if nucleoside-modified mRNA-LNP can drive the expression of type I IFNs within the dLNs to impact the induction of GC responses. We first attempted to measure local IFN-β levels by ELISA in the dLNs following injection of mRNA-LNP, as previously done for IL-6^8^. However, the levels of IFN-β were below the limit of detection when using a conventional ELISA approach (data not shown). Since type I IFNs are powerful immunomodulators^22,24^, even when present at low levels, we next asked whether the *in vivo* blockade of type I IFN signaling influenced the generation of Tfh cell and GC B cell responses to mRNA-LNP vaccination. To this end, we administered an interferon (alpha and beta) receptor 1 (IFNAR) blocking antibody to mice 24 hours prior to intramuscular (IM) immunization with an mRNA-LNP vaccine (**Figure 1A**). In the absence of IFNAR signaling, the generation of GC responses was hampered, as demonstrated by a decrease in the number of Tfh cells and GC B cells (**Figure 1B-C and S1A-B**). However, the frequency of both cell types remained intact. Together, these data indicate that nucleoside-modified mRNA-LNP can induce a low-level production of type I IFNs that is biologically relevant for the amplification of GC responses.

**Figure 1.**
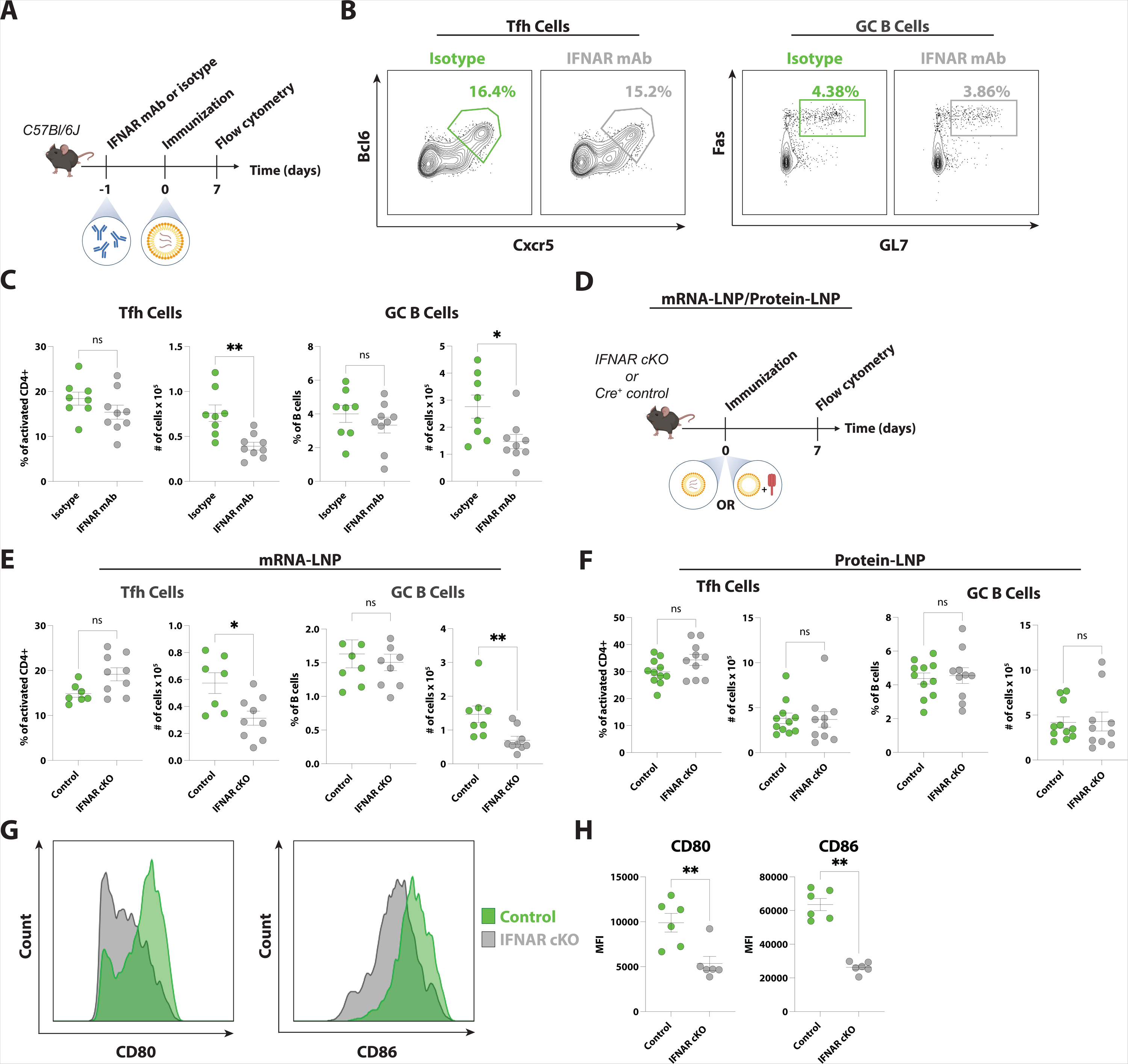
Type I IFN signaling in DCs amplifies GC responses to mRNA-LNP. **(A)** Experimental design of panels **B** and **C.** C57BL6/J mice received anti-IFNAR or isotype control monoclonal antibody (mAb) one day prior to immunization. **(B)** Representative flow cytometry of Tfh cells (**Left**; Live, B220^−^CD4^+^CD44^+^CD62L^−^ Cxcr5^+^Bcl6^+^) and GC B cells (**Right**; Live, CD19^+^CD3^−^Fas^+^GL7^+^). **(C)** Tfh cell (**Left**) and GC B cell (**Right**) frequency and absolute numbers were analyzed, as detailed in panel **B**, 7 days post-immunization. **(D)** Experimental design of panels **E** and **F.** Control (CD11c-cre^+^) or IFNAR cKO (CD11c-cre IFNAR^flox/flox^) mice were used in this study. **(E and F)** Tfh cell (**Left**) and GC B cell (**Right**) frequency and absolute numbers were analyzed, as detailed in panel **B**, 7 days post-immunization with mRNA-LNP (**E**) or protein-LNP (**F**). **(G)** Representative fluorescent intensity of CD80 and CD86 on the surface of DCs (Live, CD3^−^ CD19^−^CD11c^+^MHCII^+^) 24 hours post-immunization in C57BL6/J mice. Fluorescent intensity is displayed as count (normalized to mode). **(H)** Quantification of CD80 and CD86 expression from the experiment described in **G**. In (**A-E**), mice received a single IM immunization with 30ug of influenza virus hemagglutinin (HA) mRNA-LNP. In (**F**), mice received a single IM immunization with 30μg of recombinant HA combined with LNP. In (**G** and **H**) mice received a single IM immunization with 30ug of RBD mRNA-LNP. In (**C-F**), n = 7-11 mice per group. In (**H**), n = 6 mice per group. Data were combined from 2-3 independent experiments. Statistical analysis: unpaired two-tailed Mann-Whitney *U* test was conducted. Error bars represent SEM. *p ≤ 0.05, **p ≤ 0.01.

Next, to test if IFNAR signaling in DCs is responsible for the defective amplification of GC responses, as observed in certain models^25,28^, we immunized mice lacking IFNAR in DCs (CD11c-cre IFNAR^flox/flox^; IFNAR cKO) and littermate controls (CD11c-cre IFNAR^wt/wt^) with an mRNA-LNP vaccine (**Figure 1D**). We found that IFNAR cKO mice had normal frequencies of Tfh and GC B cells coupled to lower numbers of both GC cell populations, mirroring the outcome of our antibody blockade experiments (**Figure 1E**). Interestingly, when the IFNAR cKO mice were immunized with a protein antigen mixed with an equivalent amount of LNP (protein-LNP), the GC responses were intact (**Figure 1D and F**). To determine the mechanism of nucleoside-modified mRNA adjuvanticity in driving optimal GC responses, we next investigated whether the absence of IFNAR signaling impacts the maturation of DCs. This can be measured by the expression of the costimulatory molecules CD80 and CD86^29^. An impaired expression of costimulatory molecules by IFNAR cKO DCs could explain why we observed Tfh cell responses that were normal in frequency but reduced in numbers in IFNAR cKO mice. We immunized IFNAR cKO and control mice with mRNA-LNP and assessed the expression of CD80 and CD86 on DCs from the dLN 24 hours following immunization. DCs lacking IFNAR had a marked decrease in the expression of the costimulatory molecules CD80 and CD86 (**Figure 1G and H**). Lastly, we tested whether type I IFN signaling in DCs is important for the production of the pro-Tfh cytokine IL-6 after mRNA-LNP vaccination, as shown in certain viral infections^25^. However, a normal level of IL-6 was found in dLNs of IFNAR cKO mice in comparison to control mice (**Figure S1C**). Taken together, these data suggest that the *in vivo* sensing of nucleoside-modified mRNA contributes to the adjuvant effect of mRNA-LNP vaccines by favoring a type I IFN-dependent DC maturation/activation that, in turn, promotes optimal Tfh cell and GC responses.

### The nucleoside-modified mRNA and LNP vaccine components drive divergent CD4 T cell functional profiles

In previous work, we demonstrated that different vaccine modalities drive the generation of functionally distinct Tfh cells^9^. Specifically, mice immunized with mRNA vaccines generated Tfh cells which secreted predominantly IFN-γ (and IL-4 to a lesser extent) upon antigen restimulation, indicating a more pronounced Th1 polarization. By contrast, mice immunized with protein antigen formulated in AddaVax, an MF59-like adjuvant, mounted Tfh cell responses that were skewed toward a Th2 profile, as shown by the enrichment in IL-4-producing Tfh cells in this group. Since mRNA sensing through TLR7 can be linked to a Th1-biased response^30,31^, we hypothesized that the Tfh cell responses elicited by mRNA-LNP and LNP-adjuvanted protein might be qualitatively different (although similar in magnitude)^9^. To test this hypothesis, we immunized mice with either mRNA-LNP or antigen-matched protein-LNP. CD4 T cells from dLNs were isolated at day 7 post-immunization and restimulated with PMA/Ionomycin, prior to intracellular cytokine staining (**Figure S2A**). Activated CD4 T cells from mice immunized with mRNA-LNP were enriched in IFN-γ-producing cells, while those from mice immunized with protein-LNP had a trend for a lower frequency of IFN-γ^+^ cells along with a higher frequency of IL-4^+^ cells (**Figure 2A and B**). A more accentuated divergence was observed for Cxcr5^+^ Tfh cells (**Figure 2B and S2B**). Accordingly, the analysis of the IL-4^+^ to IFN-γ^+^ cell ratio confirmed a stronger Th1 polarization in both total CD4 and Tfh cells from mice immunized with mRNA-LNP as opposed to a more overt Th2-skewing of CD4 T cell populations in mice immunized with protein-LNP (**Figure 2C**).

**Figure 2.**
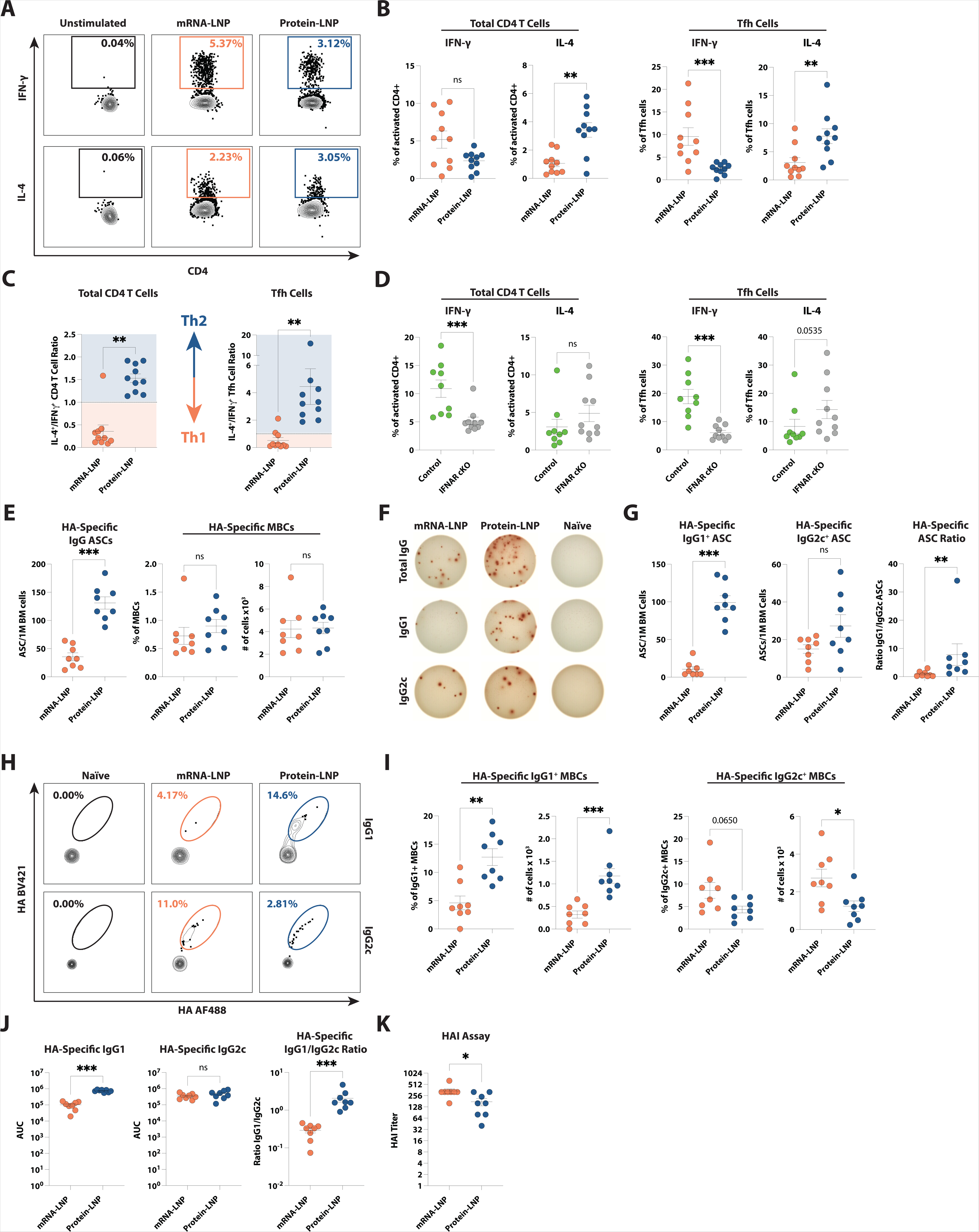
Nucleoside-modified mRNA drives a Th1-biased immune response through a type I IFN-dependent mechanism. **(A)** Representative flow cytometry analysis of IL-4 and IFN-γ expression in activated CD4 T cells (Live, CD19^−^CD4^+^CD44^+^). CD4 T cells were isolated from dLNs of C57BL/6J mice 7 days post-immunization and restimulated with PMA/Ionomycin before measurement of IL-4 and IFN-γ. **(B)** Quantification of IL-4 and IFN-γ cytokine production by total activated CD4 T cells (**Left**) and Tfh cells (**Right**; Live, CD19^−^CD4^+^CD44^+^Cxcr5^+^PD-1^+/-^) from the experiment described in **A**. **(C)** Ratio of IL-4^+^/ IFN-γ^+^ CD4 T cells (**Left**) or Tfh cells (**Right**) from panel **B**. **(D)** CD4 T cells were isolated from the dLNs of control (CD11c-cre^+^) or IFNAR cKO (CD11c-cre IFNAR^flox/flox^) mice 7 days post-immunization. Cells were stimulated prior to quantification of IL-4 and IFN-γ cytokine production by total activated CD4 T cells (**Left**) and Tfh cells (**Right**). **(E)** Antigen-specific IgG^+^ ASCs (**Left**) and MBCs (**Right**) were measured by ELIPSOT or flow cytometry, respectively, 70 days post-immunization. **(F)** Representative ASCs measured by ELIPSOT 70 days post-immunization. ASCs are represented as the number of ASCs per 1 million bone marrow cells. Corresponding to Figure 2E and G. **(G)** Quantification of antigen-specific IgG1^+^ (**Left**) and IgG2c^+^ (**Middle**) ASCs, measured as described in **F**. Ratio of IgG1^+^/IgG2c^+^ ASCs (**Right**). **(H)** Representative flow cytometry of HA-specific MBCs 70 days-post immunization, corresponding to Figure 2I. **(I)** Frequency and number of antigen-specific IgG1^+^ (**Left**) and IgG2c^+^ (**Right**) MBCs from **H**. **(J)** Antigen-specific IgG1 and IgG2c antibody level 70 days post-immunization, calculated as area under the curve (AUC). **(K)** Hemagglutination Inhibition Assay (HAI) measured 70 days post-immunization. In (**A-D**), mice received a single IM immunization with 30ug of RBD mRNA-LNP or 30μg of recombinant RBD combined with LNP, as indicated. In (**E-K**) mice received a single IM immunization with 30ug of HA mRNA-LNP or 30μg of recombinant HA combined with LNP, as indicated. In (**A-F**), n = 9-10 mice per group. In (**E-K**), n = 8 mice per group. Data were combined from 2-3 independent experiments. Statistical analysis: unpaired two-tailed Mann-Whitney *U* test was conducted. Error bars represent mean + SEM. *p ≤ 0.05, **p ≤ 0.01, ***p ≤ 0.001, ****p ≤ 0.0001.

Sensing of type I IFNs by DCs can either favor or antagonize the generation of CD4 T cell responses with a Th1 profile^32^. To determine whether type I IFN signaling in DCs in response to the nucleoside-modified mRNA-LNP vaccines was responsible for the enhanced Th1 polarization of total CD4 T cells and Tfh cells, we immunized IFNAR cKO and control mice with mRNA-LNP and assessed the polarization of CD4 T cells from dLN. We observed that the IFNAR cKO mice had a significant decrease in the frequency of IFN-γ-producing CD4 T cells and Tfh cells in comparison to the control mice, whereas the IL-4 production potential of both CD4 T cell populations was intact (**Figure 2D**). These data pointed to a model where the nucleoside-modified mRNA-driven type I IFNs act on DCs to promote Th1-polarized CD4 T cell responses.

We next interrogated the capacity of nucleoside-modified mRNA to influence additional functional properties of Tfh cells. One of the key cytokines produced by Tfh cells to support the generation of affinity-matured B cell responses is IL-21^12,13^. Hence, we evaluated IL-21 production by Tfh cells after immunization with mRNA-LNP or protein-LNP. Protein-LNP immunization was associated with a trend towards a more robust induction of IL-21 by Tfh cells than mRNA-LNP (**Figure S2C and D**), which was not dependent on type I IFN sensing by DC, as shown by the normal frequency of IL-21^+^ Tfh cells from IFNAR cKO mice after immunization with mRNA-LNP (**Figure S2D**). The finding that nucleoside-modified mRNA sensing promotes a Th1 polarization of Tfh cells along with a trend for decreased production of IL-21 prompted us to compare the quality of B cell responses driven by mRNA-LNP versus those elicited by a protein-LNP vaccine. Thus far, either an influenza PR8 hemagglutinin (HA) (HA mRNA-LNP) or SARS-CoV-2 receptor binding domain (RBD mRNA-LNP) has been used for the immunizations. Since RBD mRNA-LNP drives modest memory B cell (MBC) responses in C57BL/6 mice (data not shown), the HA mRNA-LNP was used in this study. We assessed the formation of antigen-specific antibody-secreting cells (ASC) and antigen-specific MBCs (**Figure S2E**) at day 70 post-immunization. Our data revealed an inferior capacity of the mRNA-LNP vaccine to induce antigen-specific ASCs, whereas MBC responses specific to the antigen delivered by vaccinations were comparable in the two vaccine groups (**Figure 2E and F**). In mice, the polarization of CD4 helper T cells is reflected by the isotype of antibody produced, with Th1-biased CD4 T cells driving IgG2a/c and Th2-polarized cells leading to IgG1 production^33–35^. Hence, we evaluated the isotype of antibodies produced by antigen-specific ASCs and MBCs in response to mRNA-LNP and protein-LNP vaccines. In agreement with the more robust Th1 polarization instructed by the nucleoside-modified mRNA, the mRNA-LNP vaccines were limited in their ability to promote the generation of IgG1 ASCs (**Figure 2F and G**) and MBCs (**Figure 2H and I**), whereas their capacity to produce IgG2c ASCs and MBCs was intact or increased, respectively, in comparison to the protein-LNP vaccine. In line with these data, mRNA-LNP-immunized mice presented with comparable IgG2c and significantly lower IgG1 and IgG1/IgG2c antibody ratios when compared to the protein-LNP-immunized mice (**Figure 2J**). Of note, despite the overall lower production of ASC, immunization with mRNA-LNP yielded superior HAI titers (**Figure 2K**), suggesting that even if lower in numbers, antibody responses elicited by mRNA-LNP might be qualitatively superior to those induced by protein-LNP.

Altogether, these data indicate that the nucleoside-modified mRNA and LNP components of mRNA vaccines possess distinct adjuvant properties, with the nucleoside-modified mRNA driving pronounced Th1-polarization of total CD4 and Tfh cell responses through a type I IFN-dependent mechanism and LNP driving an enrichment in Th2 polarization and more abundant antigen-specific antibody-secreting cells.

### LNP-containing vaccines instruct distinct transcriptional states in DCs

We next sought to evaluate the impact of the nucleoside-modified mRNA and LNP vaccine components on the transcriptional program of DCs, with a focus on pathways and specific signals associated with the induction of Tfh cells. To this aim, we deployed the LIPSTIC system to probe the transcriptional profile of DCs in relation to their antigen-presenting status^36,37^. The LIPSTIC technology relies on proximity-dependent labeling using the Staphylococcus aureus transpeptidase sortase A (SrtA), which can covalently transfer a biotinylated substrate from TCR transgenic OT-II cells expressing SrtA bound to CD40L (CD40L^SrtA^) to acceptor DCs expressing a five N-terminal glycine residue tag on CD40 (CD40^G5^) upon antigen presentation. We designed mRNA and recombinant protein constructs where the RBD of the SARS-CoV-2 spike protein was fused via a GS linker to the chicken ovalbumin peptide sequence recognized by OT-II CD4 T cells (aa 318-340, referred to as RBD-OVA) (**Figure 3A and S3A and B**). OT-II-CD40L^SrtA^ cells were transferred into CD40^G5^ mice prior to IM immunization with either RBD-OVA mRNA-LNP, recombinant RBD-OVA with LNP (protein-LNP), or recombinant RBD-OVA with AddaVax (protein-AddaVax; used as a comparator in previous studies^8,9^). A biotin-conjugated LPETG substrate was then administered 22 hours post-immunization to track the DCs that primed the OTII CD4 T cells (biotin^+^ DCs). Total DCs from dLNs were isolated 24 hours post-immunization and the transcriptional profiles of the biotin-labeled and unlabeled DCs were determined via single-cell RNA sequencing. Of note, the principal component analysis (PCA) of total (biotin^+^ and biotin^−^) DCs revealed that the largest contributor to the variance was the presence of LNP in the vaccine formulation (PC1, over 70%), whereas the presence of nucleoside-modified mRNA accounted for only 15% of the variance in PC2 (**Figure 3B**). This finding indicated that LNP were the major drivers for the transcriptional changes observed between DCs from mice immunized with LNP-adjuvanted or AddaVax-adjuvanted vaccines. A UMAP visualization of the data before integration also confirmed that DCs from mice immunized with protein-AddaVax displayed discrete and largely non-overlapping clusters in comparison to the DCs from animals receiving LNP-adjuvanted vaccines (**Figure S3C**). A dot plot of activation markers along with molecules involved in T cell priming and migration/chemotaxis showed differential expression in the LNP-adjuvanted groups compared to the AddaVax group (**Figure 3C**). Furthermore, this analysis verified our earlier findings that the nucleoside-modified mRNA drives an upregulation of the costimulatory molecules CD80/CD86 (**Figure 1G and H**). When evaluating the distribution of the biotin^+^ DCs (indicative of antigen presentation to OT-II cells) before integration, we observed that the biotin signal was spread out across most clusters in LNP-adjuvanted groups (**Figure S3D**). Different from what was previously observed with a weaker adjuvant (Alum) or tumor-presenting DCs^36,37^, this broad distribution of biotin labeling suggests that strong adjuvants, like LNP, drive an activating effect on most DCs within the dLNs, dwarfing the effect driven by cognate T cell help on DCs.

**Figure 3.**
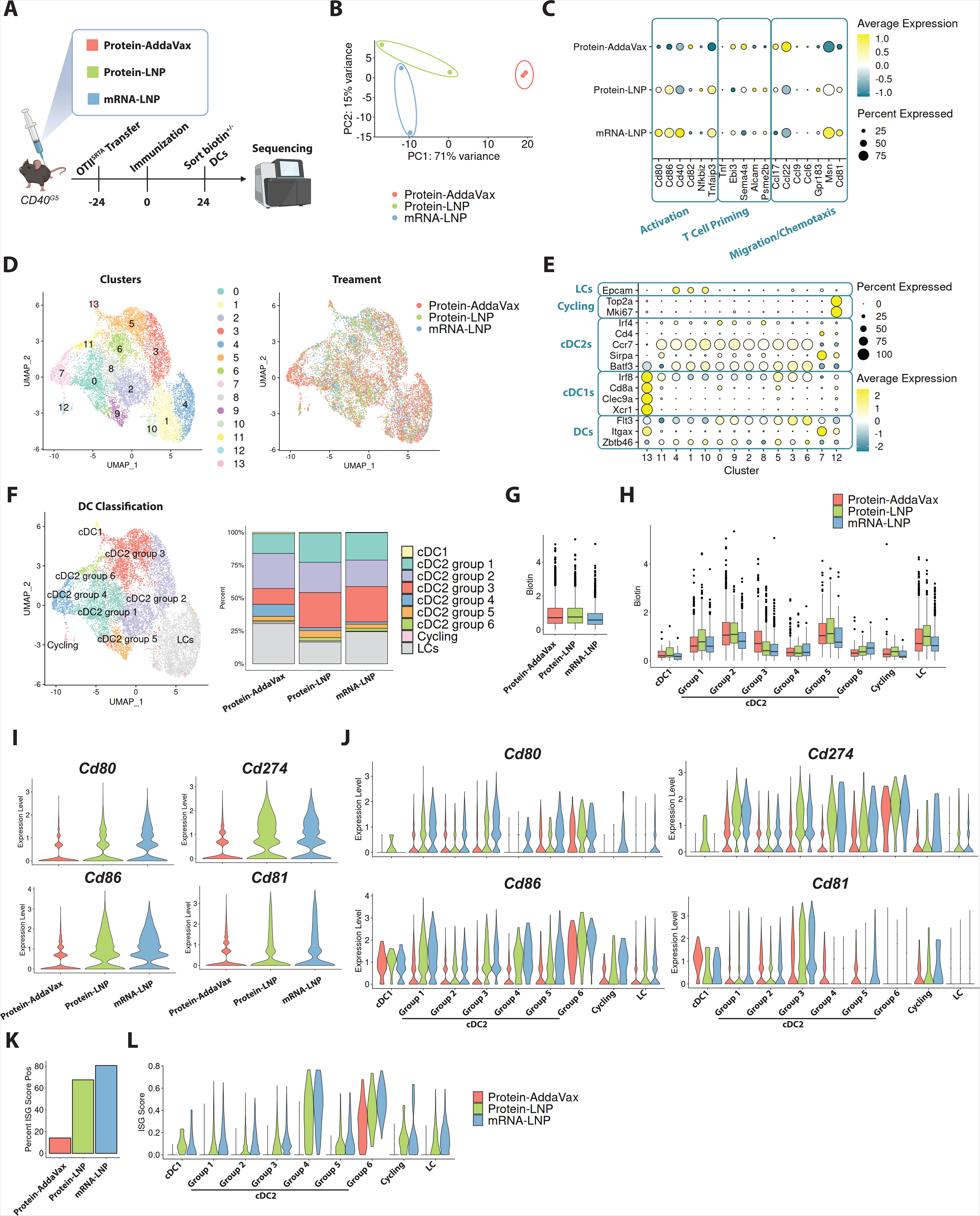
LNP-adjuvanted vaccines drive distinct gene expression profiles. **(A)** LIPSTIC Experimental setup. OT-II T cells were adoptively transferred 24 hours prior to immunization with either mRNA-LNP, protein-LNP, or protein-AddaVax. 24 hours post-immunization biotin^+^ and biotin^−^ DCs were sorted for sequencing. **(B)** PCA of LIPSTIC single-cell RNA sequencing. Each point represents an individual mouse. **(C)** Dot plot indicating the expression level of activation, T cell priming, and migration/chemotactic marker profiles driven by the treatment group. **(D)** CCA-integrated UMAP clustering displayed as unsupervised clusters (**Left**) or by treatment group (**Right**). **(E)** Dot plot displaying the gene list used to define clusters in (**D)**. **(F)** DC classification of UMAP clusters based on gene expression profiles in **E** (**Left**), also displayed as frequency of each treatment group **(Right)**. (**G and H**) Frequency of biotin labeling based on treatment group (**G**) and on the different DC clusters (**H**). (**I and J**) Expression of the indicated genes (*Cd80*, *Cd86*, *Cd274* and *Cd81*) in each treatment group (**I**) and on the different DC clusters (**J**). **(K)** Interferon stimulated gene (ISG) signature expression score in each treatment group. **(L)** ISG signature in each DC group from **F,** separated by immunization condition. Mice were immunized with either 30μg of RBD-OVA mRNA-LNP, 30μg recombinant RBD-OVA combined with LNP, or 30μg recombinant RBD-OVA combined with AddaVax; n= 2 mice per group.

Several subtypes of DCs exist which are endowed with distinct T cell priming and activation properties^38^. While type 1 conventional DCs (cDC1) are effective at cross-presentation to CD8 T cells, type 2 conventional DCs (cDC2) are specialized in antigen presentation to CD4 T cells and induction of Tfh cells ^29,39^. Canonical Correlation Analysis (CCA) in Seurat^40^ was performed to integrate transcriptomic data from the three immunization conditions and generated clusters that were for the most part evenly represented in each immunization condition (**Figure 3D**). A curated gene list (**Figure 3E**) was then used to determine the types of DCs that resided in each cluster, leading to the definition of cDC1, cDC2, Langerhans cells (LCs), and cycling cells. Upon renaming the UMAP clusters accordingly (**Figure 3F**), the vast majority of the cells were cDC2s of differing transcriptional states. We therefore assigned the cDC2s to 6 different groups based on their top differentially expressed genes (**Figure S3E**). Next, we sought to determine if the distribution of biotin differed within the DC clusters identified. The biotin signal was similarly distributed across the protein-adjuvanted vaccine groups, with a modest decrease in biotin enrichment in the mRNA-LNP group (**Figure 3G**). As anticipated, the cells with the highest biotin expression did not fall in the cDC1 cluster and were enriched in the LC cluster and selected cDC2 groups (Groups 1,2 and 5; **Figure 3H**). Additionally, a GSEA approach confirmed the enrichment of a previously published LIPSTIC gene signature^37^ in biotin^+^ cells, further validating the antigen-presenting nature of the biotin^+^ DC (**Figure S3F and G**).

Since we observed that LNP-containing vaccines drove gene signatures of activation and migration on DCs (**Figure 3C**), we next asked whether the expression of these signature genes was restricted to specific DC populations. When looking at the expression of the costimulatory molecules CD80 and CD86 in each cluster, broken up by immunization group (**Figure 3I and J**), we found a marked increase in the expression of these molecules on cDC2s from the LNP-containing groups, when compared to the AddaVax group. Among the three cDC2 groups with the highest expression of *Cd80* and *Cd86* (groups 1, 4 and 6) in mice immunized with mRNA-LNP or protein-LNP, only one was enriched in biotin expression (group 1, **Figure 3H and J**), suggesting that some groups of DCs that did not yet present antigen might have gained enhanced maturation as a result of the immunostimulatory activity of the nucleoside-modified mRNA and/or the LNP. Regulators of migration were also more robustly driven by LNP-containing vaccines in comparison to AddaVax (**Figure 3I and J** and **Figure S3H and I**), as indicated by a noticeable induction of *Cd81* (a regulator of DC motility^41^), *Cd274* (the gene encoding for PD-L1, which is also important for the migratory capacity of DCs^42^), and *Cxcr5* (a chemokine receptor important for the migration of DCs toward B cell follicles^43^), in many cDC2 groups. Finally, given the connection between type I IFNs and nucleoside-modified mRNA sensing (**Figure 1**), we asked whether an enrichment in an interferon-stimulated gene (ISG) signature was specifically present in the animals immunized with mRNA-LNP. To this aim, we utilized an ISG signature consisting of the top 20 differentially expressed genes in DCs when IFN-α was applied *in vitro* as described in^44^. We found that the ISG signature was enriched in the mRNA-LNP immunization condition, and less so in the protein-LNP immunization group, when compared to the protein-AddaVax immunization group (**Figure 3K and L**). When stratified by DC subtypes, we found an enrichment of the ISG signature in biotin^lo^ cDC2 groups 4 and 6, suggesting a direct induction of type I IFN responses directed by the nucleoside-modified mRNA (and to a lesser extent by the LNP). Taken together, these data support the notion that the LNP component of mRNA vaccines is a powerful immunostimulator that drives activation and migration programs in cDC2s, whereas the nucleoside-modified mRNA component appears to have a less notable contribution to the transcriptional changes driven by mRNA-LNP vaccines.

### LNP drive the induction of a pro-Tfh cell program in cDC2s and guide Tfh cell differentiation via an Ebi2-oxysterol axis

We next sought to determine if LNP and/or the nucleoside-modified mRNA were capable of efficiently inducing a “pro-Tfh” program in cDC2s. Since groups 4 and 6 cDC2s expressed high levels of activation, migration and ISG signatures and were biotin^lo^ (hence the transcriptional changes did not result from T cell help following antigen presentation), we asked what additional pro-Tfh genes were enriched in these groups in response to LNP adjuvanted vaccines. Previous work has demonstrated that DCs positioned at T-B borders can promote the differentiation of Tfh cells by interacting with Ebi2-expressing activated CD4 T cells and producing a soluble form of CD25 that quenches the cytokine IL-2, a potent inhibitor of Tfh cell differentiation^15–17^. Interestingly, we found that the expression of *Il2ra* (the gene encoding for CD25) and *Gpr183* (the gene encoding for Ebi2) were both highly enriched in cluster 6, whereas cluster 4 displayed heightened expression of *Grp183* but not of *Il2ra*, suggesting that these two clusters might have discrete pro-Tfh properties (**Figure 4A**). Moreover, both genes were expressed in DCs from mice immunized with LNP-containing vaccines (**Figure 4B**), further indicating that LNP inherently drive a “pro-Tfh” program in DCs.

**Figure 4.**
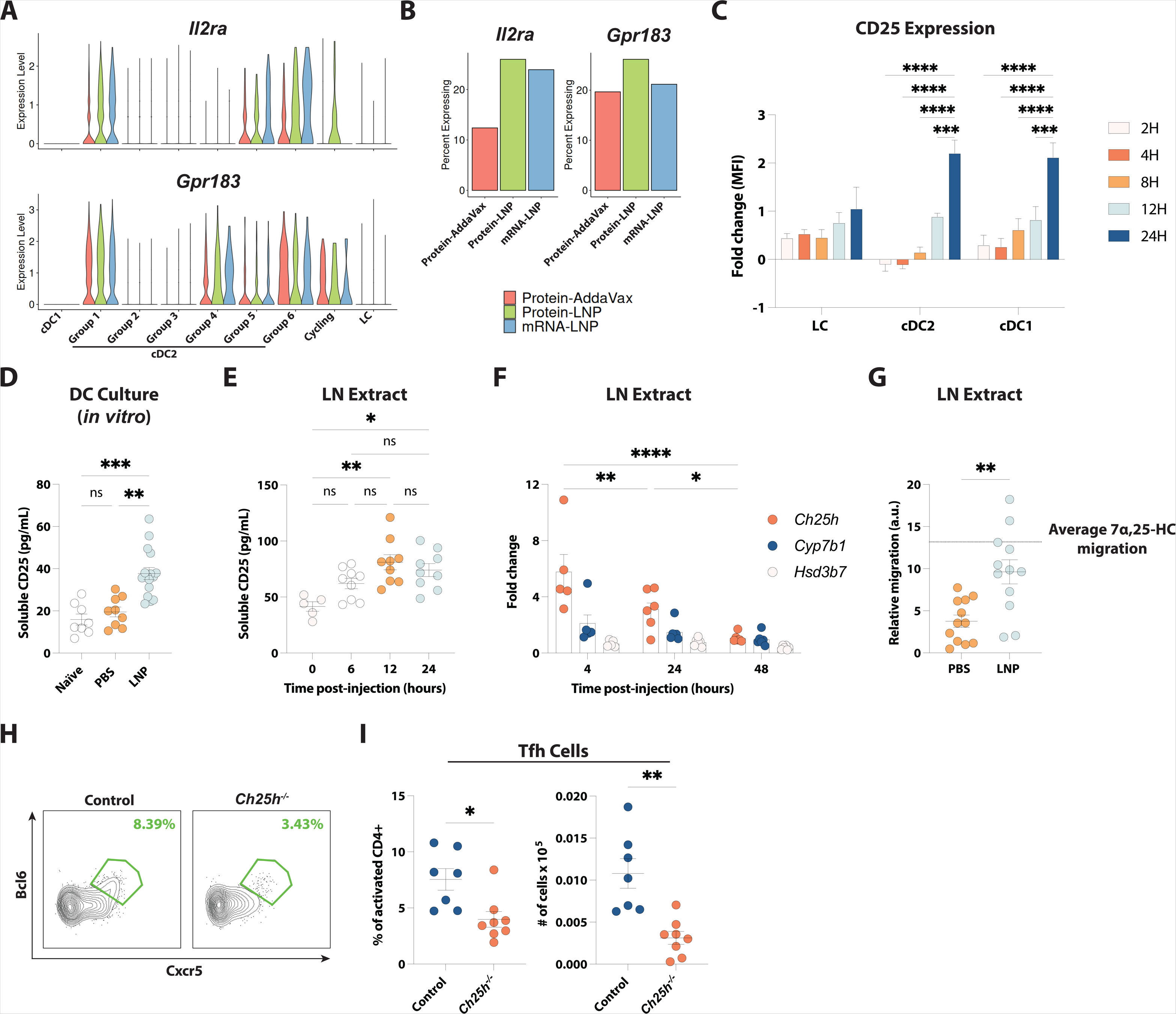
LNP-adjuvanted vaccines drive an enrichment of CD25 and Ebi2 expression in DCs. **(A)** Violin plots of Il2ra and Gpr183 in DC clusters from Figure 3F, separated by immunization condition. **(B)** Expression of *Il2ra* and *Gpr183* expression in each immunization condition from RNA-sequencing data described in Figure 3. **(C)** Surface expression of CD25 on DC subsets measured by flow cytometry at the indicated time points post-immunization, represented as fold-change over baseline expression (0H). **(D)** Soluble CD25 produced *in vitro* was measured by ELISA. LN DCs were isolated 24 hours post-injection of LNP or PBS and cultured *in vitro* for 24 hours before analysis of CD25 in the culture supernatants. **(E)** Soluble CD25 produced *in vivo* was measured by ELISA. Whole LN extracts were taken at the indicated time points post-injection with LNP before analysis of CD25 in the extract supernatants. **(F)** qPCR analysis of cholesterol-converting enzymes in LN extracts obtained at the indicated time points after LNP injection. Data is displayed as fold change versus naïve mice. **(G)** M12 cell migration in response to lipid extracts from dLNs of mice injected with PBS or LNP 24 hours prior. Dotted line denotes the average migration to 7α,25-HC. **(H)** Representative flow cytometry of Tfh cells in *Ch25h^−/−^* versus control mice 7 days post-immunization. **(I)** Quantification of Tfh cells frequency and number in *Ch25h^−/−^* and control mice from **H**. In (**C**), mice received a single IM immunization containing 20 μg of eGFP mRNA-LNP-DiI. In (**D-F**) mice received a single IM injection of LNP or PBS, as indicated. In (**G** and **H-I**) mice received a single IM immunization with 30 μg of RBD mRNA-LNP. Statistical analysis: (**C** and **F**) Two-way ANOVA was performed with Tukey’s correction for multiple comparisons. Only sample groups were compared versus all time points. (**D** and **E**) One-way ANOVA Kruskal-Wallis test was performed with Dunn’s correction for multiple comparisons. (**G** and **I**) unpaired two-tailed Mann-Whitney *U* test was conducted. Error bars represent mean + SEM. *p ≤ 0.05, **p ≤ 0.01, ***p ≤ 0.001, ****p ≤ 0.0001.

To gain further insights into how LNP shape the pro-Tfh program in cDC2s, we evaluated the induction of CD25 in DCs upon *in vivo* administration of mRNA-LNP. We observed an increase in the surface level of CD25 on all DC types, which peaked 24 hours after the injection of mRNA-LNP and was more overt in cDC1 and cDC2 than in LCs (**Figure 4C and S4A**). Furthermore, we demonstrated that the LNP vaccine component was sufficient to promote the production of the soluble form of CD25 by DCs, as shown by the detection of CD25 in the supernatant from DCs isolated from dLNs (**Figure 4D**) or from total LN extract (**Figure 4E**) following *in vivo* exposure to LNP. These data suggested that, as previously demonstrated in other immunization settings, LNP might favor Tfh cell differentiation by promoting the quenching of IL-2 via CD25 expression on DCs.

Ebi2 expression is critical for the proper positioning of DCs and CD4 T cells at the T-B borders of secondary lymphoid organs^13,17,45,46^. Hence, we investigated whether, in addition to driving *Gpr183* expression, LNP were capable of promoting the production of the Ebi2 ligand, 7α, 25-dihydroxycholesterol (7α, 25-HC)^47,48^. Firstly, we measured the capacity of LNP to induce the expression of the genes encoding the two enzymes required for the conversion of cholesterol into 7α, 25-HC^47^. We found that the *in vivo* administration of LNP led to upregulated expression of *Ch25h* and *Cyp7b*, which are both required for the conversion of cholesterol into 7α, 25-HC, but not *Hsd3b7*, which further metabolizes this oxysterol into a derivative that can no longer be recognized by Ebi2^47,48^ (**Figure 4F**). Next, to determine if the increased expression of *Ch25h* and *Cyp7b1* driven by LNP was associated with an increase in Ebi2 ligands, we adopted a previously described *in vitro* bioassay^49^. In this assay, the relative migration of an Ebi2-expressing cell line compared to a control identical cell line was assessed in response to the lipid extract from dLNs of LNP- or PBS-injected-mice. We observed that the injection of LNP increased the production of Ebi2 ligands *in vivo* (**Figure 4G**), as measured by the more robust migration in response to the lipid extract from the dLN of the LNP-injected mice in comparison to the PBS-treated group. To ascertain if the heightened production of Ebi2 ligands driven by LNP is relevant to Tfh cell differentiation *in vivo*, we immunized mice lacking CH25H (*Ch25h^−/−^*) with mRNA-LNP and assessed Tfh cell responses 7 days later. Our experiment revealed a decrease in the relative and absolute numbers of Tfh cell responses in *Ch25h^−/−^* mice compared with control mice (**Figure 4H and I**). Since CH25H is involved in the production of the Ebi2 ligand 7α, 25-HC, our data indicate an essential role of LNP-promoted 7α, 25-HC production in driving effective GC responses following mRNA-LNP immunization.

### Conventional dendritic cells are efficient at internalizing mRNA-LNP

Since we demonstrated that LNP can drive the induction of *Il2ra* and *Gpr183* in cDC2, we asked whether a direct uptake of LNP was associated with the capacity to instruct a pro-Tfh program in these cells. To address this question, we developed custom mRNA-LNP in which mRNAs encoding an enhanced green fluorescent protein (eGFP) were encapsulated in LNP labeled with a fluorescent lipophilic dye (1,1’-Dioctadecyl-3,3,3’,3’-Tetramethylindocarbocyanine Perchlorate (referred to as DiI); **Figure 5A**). DiI signal allows for tracking of LNP binding and/or uptake whereas eGFP positivity provides a measure of mRNA translation into protein. cDC2s from the dLNs were analyzed 24 hours after the injection of mRNA-LNP using the gating strategy described in **Figure S5A.** We observed an increased CD25 expression in the cDC2s that had bound/internalized LNP (DiI^+^), regardless of the mRNA translational status (eGFP^−^ and eGFP^+^) (**Figure 5B**). Similarly, the LNP-driven induction of Cxcr5 was more prominent in DiI^+^ cDC2 populations compared to DiI^−^ cDC2s (**Figure 5C**), suggesting that the immunostimulatory effect of the LNP is predominantly restricted to the cells that have bound/internalized LNP.

**Figure 5.**
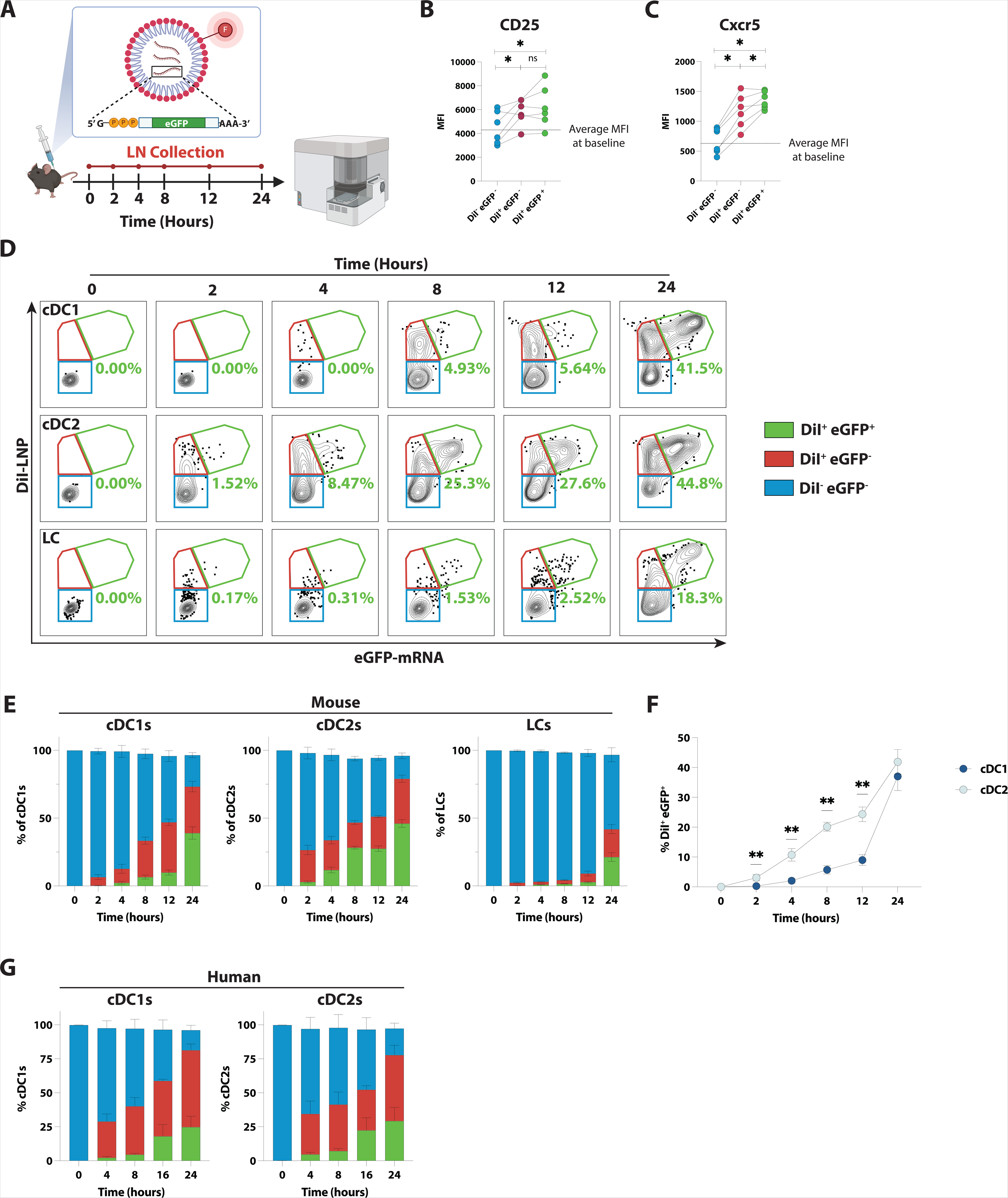
Conventional dendritic cells are efficient at internalizing LNP and translating mRNA. **(A)** Experimental scheme. Mice were injected with an eGFP-encoding mRNA encapsulated in a DiI-labeled LNP (eGFP mRNA-LNP-DiI). Following immunization, dLNs were collected at the indicated time points, and lymphocyte populations were analyzed by flow cytometry. **(B and C)** CD25 **(B)** and Cxcr5 **(C)** expression in DiI^−^ eGFP^−^, DiI^+^ eGFP^−^ and DiI^+^ eGFP^+^ cDC2s (CD11c^+^MHCII^+^CD24^+^EpCAM^−^CD103^−^CD11b^+^), 24 hours post eGFP mRNA-LNP-DiI injection. The dotted line indicates the average mean fluorescence intensity (MFI) in cDC2s of naïve (0H) mice. **(D)** Representative flow cytometry of LNP binding/uptake (DiI^+^) and mRNA translation (eGFP^+^) in cDC1s (CD11c^+^MHCII^+^CD24^+^EpCAM^−^CD103^+^CD11b^−^), cDC2s (CD11c^+^MHCII^+^ CD24^+^EpCAM^−^CD103^−^CD11b^+^), or LCs (CD11c^+^MHCII^+^CD24^+^EpCAM^+^) from dLN of mice at different time points post eGFP mRNA-LNP-DiI injection. **(E)** Quantification of **D**. Each vertical bar represents 100% of the indicated DC subtypes and color represents the portion of cells that are DiI^+^ eGFP^+^ (green), DiI^+^ eGFP^−^ (red), and DiI^−^ eGFP^−^ (blue) at each post-injection time point. **(F)** Frequency of DiI^+^ eGFP^+^ cDC1s and cDC2s at the indicated time points post-injection. **(G)** Quantification of binding/uptake and translation of eGFP mRNA-LNP-DiI in human cells in vitro. cDC1s (Live, CD3^−^CD19^−^CD20^−^CD11c^+^HLA-DR^+^BDCA-3^+^BDCA1^−^) and cDC2s (Live, CD3^−^CD19^−^CD20^−^CD11c^+^HLA-DR^+^BDCA-3^−^BDCA1^+^) were isolated from healthy donor PBMCs. In (**B-F**) mice were immunized with 20μg of eGFP mRNA-LNP-DiI. For (**B-G**), n=6 per time point. Data is combined from two independent experiments. Statistical analysis: In (**B**) and (**C**), a one-tailed paired Wilcoxon test was performed. In (**F**) an unpaired Mann Whitney U test was performed with a two-stage Benjamini, Krieger and Yekutieli FDR of 1% to correct for multiple comparisons. Error bars represent mean + SEM. *p ≤ 0.05, **p ≤ 0.01.

Next, we deployed the eGFP mRNA-LNP-DiI to broadly profile the APCs present in dLNs that are capable of LNP uptake and mRNA translation. This mRNA-LNP was used in conjunction with a spectral flow cytometry panel which allowed us to discriminate cDC1, cDC2, Langerhans cells (LCs), plasmacytoid DCs (pDCs), macrophages, and inflammatory monocytes (**Figure S5A**). eGFP mRNA-LNP-DiI were injected intramuscularly, and the mRNA-LNP processing capacity of different cell types was assessed in the dLNs between 4- and 24 hours post-injection. We found that innate cell populations were capable of processing mRNA-LNP to varying degrees (**Figure S5B**), with cDCs being the most efficient at binding/internalizing LNP (DiI^+^) and translating the mRNA (eGFP^+^) and macrophages and pDCs having more modest processing capability. Interestingly, we reported a wave of DiI^+^ eGFP^+^ monocytes infiltrating into dLNs 4 hours after immunization that was largely resolved by 24 hours (**Figure S5B and C**). To further refine the kinetics of LNP binding/internalization and mRNA translation by DCs after vaccination, we injected mice intramuscularly with fluorescent mRNA-LNP and analyzed dLNs by flow cytometry after 2-, 4-, 8-, 12- and 24-hours (**Figure 5A and S5D**). In line with our previous experiment, all DC subtypes were capable of binding/internalizing LNP and translating the mRNA to some extent (**Figure 5D**). Of note, cDC2s were the fastest DC type to bind/internalize mRNA-LNP, as they began binding/internalizing LNP by 2 hours and translating mRNA by 4 hours post-immunization (**Figure 5D-F**). Despite the difference in DiI^+^ eGFP^+^ cells at early time points (**Figure 5F**), 24 hours after mRNA-LNP injection cDC1s and cDC2s displayed equivalent percentages of DiI^+^ eGFP^+^ cells. LC did not process mRNA-LNP as efficiently as cDCs. To determine if human innate cells have a comparable ability to internalize mRNA-LNP, we incubated peripheral blood mononuclear cells from 6 different healthy donors with fluorescent mRNA-LNP and analyzed processing by cDC1s and cDC2s after 4-, 8-, 16- and 24-hours of *in vitro* culture. In line with our results in mice, we found that both human cDC1s and cDC2s were able to bind/internalize mRNA-LNP within 24 hours (**Figure 5G and S5E**). It is worth mentioning that the ability of DCs from most healthy donors to translate the mRNA *in vitro* was much lower than what was seen in mice *in vivo* (**Figure S5F**). Possible explanations for this discrepancy could be the variability in mRNA-LNP processing by different subjects or increased toxicity of mRNA-LNP in an *in vitro* culture system.

Overall, our data underscore that while multiple types of innate cells can internalize LNP and translate the mRNA, cDC2 are the fastest and among the most efficient cells at processing mRNA-LNP *in vivo*.

### mRNA-LNP uptake by cDCs takes place predominantly in dLNs

The finding that most cDC2 in the dLN have internalized LNP and translated the mRNA 24 hours after the intramuscular injection of mRNA-LNP (**Figure 5**) prompted us to investigate the location of mRNA-LNP uptake. Studies in both rhesus macaques and mice have shown the rapid infiltration of various APC types, such as DCs and monocytes into the injection site upon nanoparticle-based intramuscular immunization^50–52^. In light of these data, a model was proposed where antigen uptake by migratory DCs in the muscle is followed by relocation of the DCs to the dLNs, where the DCs either directly present antigen to T cells or transfer it to LN-resident DCs for subsequent antigen presentation ^53,54^. To determine the validity of this model, we first asked if the mRNA-LNP-processing DCs found in the dLNs had a migratory or resident phenotype. Classically, migratory DCs have been characterized based on the expression level of MHC-II, with CD11c^+^ MHC-II^hi^ cells being considered migratory, while CD11c^+^ MHC-II^int^ cells are defined as resident^38,55^. Indeed, this approach defines two distinct populations in steady-state LNs (**Figure 6A**). However, upon IM injection of mRNA-LNP, MHC-II and CD11c expression both increased on all dLN DCs, and within 8 hours post mRNA-LNP administration it was no longer possible to discriminate between resident and migratory DCs.

**Figure 6.**
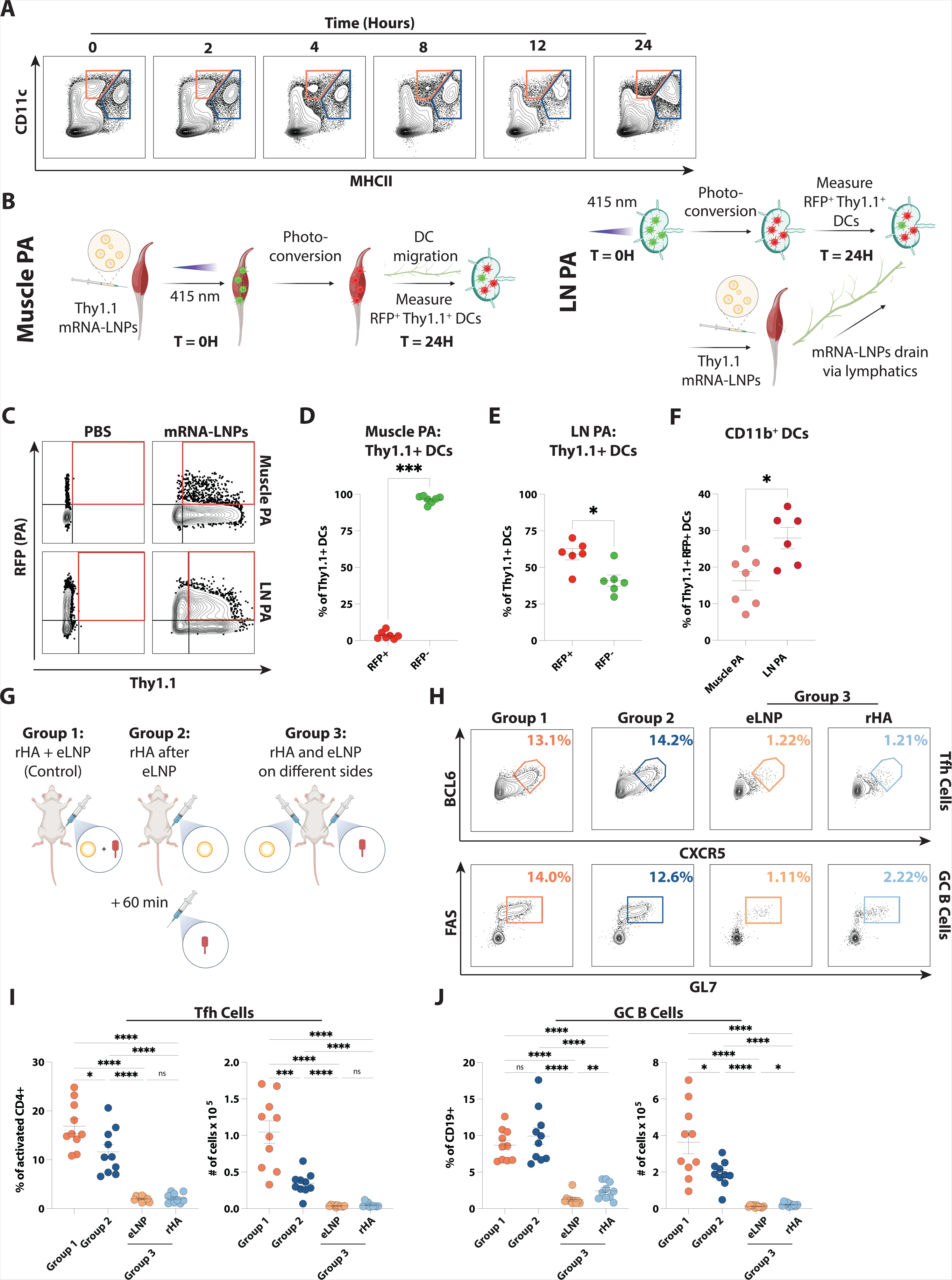
The uptake of mRNA-LNP by DCs occurs locally in draining lymph nodes. **(A)** Representative expression of MHCII and CD11c at the indicated time points post-immunization with mRNA-LNP assessed by flow cytometry. Cells are pre-gated as Live, TCRβ^−^CD19^−^. **(B)** Schematic of photoactivation experiments. Muscle photoactivation (PA) (**Left**) and LN PA (**Right**). **(C)** Representative expression of RFP (indicating photoactivation) and Thy1.1 (indicating mRNA processing) by DCs from draining LNs 24 hours after mRNA-LNP injection. Muscle PA (**Top**) and LN PA (**Bottom**) experiments are shown. **(D and E)** Quantification of RFP^+^ and RFP^−^ Thy1.1^+^ DCs in the draining LNs 24 hours after immunization in muscle PA (**D**) and LN PA (**E**) experiments. Photoactivated cells are represented as red circles while non-photoactivated cells are represented as green circles. **(F)** Frequency of CD11b^+^ (cDC2) from **D** and **E**. **(G)** Outline of immunization groups for **H**, **I** and **J**. **(H)** Representative frequencies of Tfh cells (Live, B220^−^CD4^+^CD44^+^CD62L^−^Cxcr5^+^Bcl6^+^) and GC B cells (Live, CD19^+^CD3^−^Fas^+^GL7^+^) from Groups 1, 2 and 3 (as described in **G**) measured by flow cytometry. **(I)** Quantification of frequency and number of Tfh cells 7 days post-immunization. **(J)** Quantification of frequency and number of GC B cells 7 days post-immunization. In (**A**) mice were injected with 20 μg of eGFP mRNA-LNP-DiI. In (**C-F**) mice were injected with 30 μg Thy1.1 mRNA-LNP. In (**H-J**) mice were immunized as described with either 30 μg of HA mRNA-LNP or 30 μg rHA mixed with LNP. In **A**, **C**, **D**, **E** and **F**, n = 6 mice per group combined from 2 experiments. In **G**, **H**, **I** and **J**, n = 10 mice per group combined from 2 experiments. Statistical analysis: In panels **D**, **E**, **F**, **I** and **J** an Unpaired two-tailed Mann-Whitney *U* test was performed. Error bars represent mean + SEM. *p ≤ 0.05, **p ≤ 0.01, ***p ≤ 0.001, ****p ≤ 0.0001.

To circumvent this issue, we utilized the Kikume Green-Red (KikGR) photoactivation (PA) system^56^ to specifically label muscle DCs and track their migration to the dLN in conjunction with their mRNA-LNP processing capacity (**Figure 6B**). In KikGR mice, cells express a green fluorescent protein (GFP) that is photoconverted into a red fluorescent protein (RFP) upon exposure to short-wavelength light. We first determined the efficiency of PA upon light exposure of the gastrocnemius muscle for 2 or 3 minutes. On average, about 20% of CD45^+^ cells in the muscle were RFP^+^ after 2 minutes of light exposure, while a higher percentage, about 40%, of DCs were RFP^+^ at this time (**Figure S6A and B**). Additionally, no difference was present in the frequency of PA^+^ cells in the muscle with different durations of exposure to light. Thus, the shorter 2-minute interval of light exposure was used for the following experiments.

Next, we sought to determine if muscle DC frequency changed over time following intramuscular immunization, to identify the optimal time to photoactivate the muscle. We found that the injection of mRNA-LNP promoted a rapid infiltration of CD45^+^ cells that did not express DC markers, resulting in decreased DC frequencies within 4 hours of injection (**Figure S6C**). Based on this finding, in the following experiments, the muscle was photoactivated immediately following mRNA-LNP injection, when DC frequency was highest. To track the uptake/transcription of mRNA-LNP, we designed an mRNA-LNP that encoded the Thymus cell antigen 1.1 (Thy1.1) (**Figure 6B and C**). KikGR mice were injected with Thy1.1 mRNA-LNP in the gastrocnemius muscle, which was subsequently photoactivated near the injection site. The dLNs were analyzed 24 hours following injection, revealing that only ~3.6% of the Thy1.1^+^ DCs in the dLN were RFP^+^, whereas 96.4% of Thy1.1^+^ DCs were RFP^−^. This indicated that a vast majority of mRNA-LNP-processing DCs were not in the muscle at the time of injection (**Figure 6D**). Additionally, the frequency of the Thy1.1^+^ RFP^+^ DCs in dLNs did not significantly increase in comparison to PBS-injected mice (**Figure S6D**), emphasizing that these cells comprise a minority of DCs within the dLNs.

To further understand whether mRNA-LNP uptake by DCs predominantly takes place locally in the dLN, we performed an additional set of experiments where we photoactivated the dLN directly before intramuscular injection of mRNA-LNP (**Figure 6B and C**). If mRNA-LNP reach dLNs through the afferent lymphatics and are then internalized by DCs that were already in the dLN before the immunization, then we should observe Thy1.1^+^ RFP^+^ DCs. Interestingly, we found that most Thy1.1^+^ DCs were RFP^+^ (**Figure 6E and S6E**), indicating that these DCs were present in dLNs at the time of immunization. Additionally, we found that roughly 15% of the Thy1.1^+^ RFP^+^ DCs were cDC2s when the muscle was photoactivated, however, this frequency doubled when photoactivating the dLNs (**Figure 6F**). The Thy1.1^+^ RFP^−^ DCs were likely DCs that were either not efficiently photoactivated or trafficked to the dLN as a result of the increased chemoattractant production driven by LNP.

In sum, the photoactivation experiments suggest that most dLN cDC2 internalize mRNA-LNP *in situ* rather than migrating from the muscle upon mRNA-LNP uptake.

### LNP-adjuvanted responses are restricted to the dLNs

A high dose of mRNA-LNP was used throughout this study, raising the possibility that an excess of mRNA-LNP could result in a systemic spread of the vaccine. To rule out this possibility, we did an experiment where one group of mice was immunized IM with a recombinant protein antigen (influenza A virus recombinant hemagglutinin, rHA) mixed with LNP (Group 1), a second group of mice received first LNP IM then an hour later the protein antigen on the ipsilateral side (Group 2), and a third group received the protein antigen in one leg and LNP in the contralateral leg at the same time (Group 3; **Figure 6G**). We reasoned that if the LNP spread systemically, group 3 should present GC responses at least in the LNs draining the rHA. We analyzed the induction of Tfh cells and GC B cells in the dLNs 7 days post-immunization by flow cytometry. Importantly, we did not find detectable Tfh and GC B cell responses in the dLNs draining either injection sites in Group 3, demonstrating that the LNP do not act as systemic adjuvants even when utilized at high doses (**Figure 6H-J**). Interestingly though, the analysis of Group 2 revealed that LNP do not need to be co-formulated with rHA to be affective adjuvants, as the mice that received the protein antigen one hour after the LNP presented similar (albeit slightly decreased) GC response, suggesting that the LNP and protein immunogen do not need to be physically coupled.

Overall, our findings imply that, even at high doses, LNP exert their adjuvant activity locally in vaccine-draining lymph nodes and, in the setting of protein subunit vaccination, they act in conjunction with the protein antigen.

## DISCUSSION

mRNA vaccines have been instrumental in the fight against the SARS-CoV-2 pandemic, due in large part to their ability to drive potent GC responses, which ultimately result in the production of affinity-matured neutralizing antibodies^57^. A mechanistic understanding of how mRNA vaccines elicit robust Tfh cell and GC responses, which is crucial for the rational design of mRNA vaccines with improved features, is still missing. In this study, we sought to investigate the mechanisms of Tfh cell induction by mRNA vaccines and demonstrated that the nucleoside-modified mRNA and the LNP components of mRNA vaccines each possess unique, yet complementary, adjuvant activity that cooperate to instruct an efficient GC program.

Immune cells express a multitude of pattern recognition receptors (PRR) for sensing exogenous single and double-stranded RNA (ssRNA and dsRNA), making foreign RNA a highly immunogenic molecule^58^. Among these PRRs are the endosomal receptors TLR7/8, which can sense ssRNA, and TLR3, which recognizes dsRNA along with the cytosolic sensors RIG-I and MDA5^59^. RNA recognition by these receptors triggers immune responses dominated by type I IFN production and the concomitant upregulation of ISGs. Excess production of pro-inflammatory cytokines, including type I IFNs, following RNA sensing by PRRs can have negative consequences on the translational capacity of mRNA, resulting in hindered antigen production^21,60^. However, mRNA translation can be largely restored by combining nucleoside modification with cellulose purification of the *in vitro*-produced mRNA^21,60,61^. It was originally believed that the nucleoside modification combined with purification made the mRNA completely immunosilent, as suggested by the lack of a detectable production of pro-inflammatory cytokines *in vitro* and in circulation as described in seminal *in vivo* studies from Karikó and colleagues^18,21^. Our previous finding that the TLR adaptor protein MyD88 was required to drive GC responses to mRNA vaccines^8^ led us to hypothesize that the mRNA component of our vaccines, even though nucleoside modified and purified, might still be sensed by TLRs and result in a low, but biologically relevant production of type I IFNs that contribute to GC responses. In line with our hypothesis, previous studies have reported the upregulation of ISGs in human peripheral mononuclear cells following the administration of the nucleoside-modified mRNA-LNP vaccine BNT162b2^62^. Furthermore, a role for type I IFNs in the regulation of CD8 T cell responses to the BNT162b2 was reported in mice^63^. A major limitation of these studies is, however, the fact that they only evaluated the immune responses to mRNA-LNP vaccines, preventing us from discriminating whether the type I IFN production resulted from the sensing of the mRNA or the LNP vaccine components. To overcome this major hurdle, we uncoupled the sensing of the mRNA from the sensing of the LNP by comparing the immune responses in mice immunized with an mRNA-LNP or protein-LNP vaccine. Our experimental approach revealed that type I IFN sensing by DCs is important for downstream GC reactions only in response to mRNA-containing vaccines. This data demonstrates that the mRNA component of the vaccine possesses inherent adjuvant activity, which is mediated at least in part, by the production of type I IFNs. It remains to be determined if the nucleoside-modified mRNA has some residual capacity to activate TLR7/8, or if traces of dsRNA remaining after the purification process and/or secondary structures in the nucleoside-modified mRNA^64^ trigger TLR3, resulting in type I IFN production. We do, however, speculate that TLR triggering is more relevant to promoting type I IFN production than the activation of RIG-I and MDA5, as we previously reported that MAVS-deficient animals mounted largely normal GC responses to mRNA-LNP^8^.

In our study, when type I IFN signaling was ablated in DCs, only the absolute numbers but not the frequencies of Tfh cells were decreased after immunization with mRNA-LNP. These data, combined with a lower expression of costimulatory markers on IFNAR-deficient DCs, suggest that type I IFN sensing by DCs does not alter the production of signals necessary for the Tfh cell fate commitment but is important for the priming and amplification of CD4 T cell responses. On the other hand, the mRNA-driven type I IFNs indirectly impacted the functional polarization of Tfh cells (and total CD4 T cells). Indeed, while the LNP component drove Th2-biased responses, the presence of the nucleoside-modified mRNA skewed the Tfh cell responses toward a Th1 profile. Mechanistically, we showed that the mRNA-driven Th1 polarization relies on type I IFN signaling in DCs, likely promoting the production of IL-12, a strong Th1 polarizing stimulus for CD4 T cells. This outcome was not entirely anticipated, as the impact of type I IFN signaling in DCs for driving Th1 polarization appears to be context-dependent, as suggested by type I IFN signaling resulting in IL-12-driven Th1 polarization in a *Listeria monocytogenes* infection or leading to the suppression of both Th1 and Tfh responses during *Plasmodium* infection^32^. The differential Th1/Th2 polarization molded by the nucleoside-modified mRNA and the LNP reflected a broader functional diversity instructed by the two vaccine components. Overall, we found in our study that the presence of the mRNA somehow limited IL-21 production by Tfh as well as the generation of bone marrow plasma cells, and the production of antigen-specific antibodies (especially notable was the decrease in the Th2 isotype IgG1). It is worth noticing that despite the overall decrease in plasma cells and binding antibodies, HAI titers, which are a proxy for neutralizing antibodies in influenza immunization, were more efficiently produced in mice immunized with mRNA-LNP in comparison to the protein-LNP group. Overall, this finding suggests that, at least in the setting of the specific immunogen and LNP tested, a slightly more controlled GC response (with limiting help from Tfh cells) could be more favorable for the selection of protective antibodies. Additional and currently unappreciated qualitative differences in B cell helper signals provided by Tfh cells driven by the mRNA and LNP components might also be conducive to this phenomenon and will be addressed in future studies.

When comparing the transcriptional profile of DCs from mice immunized with a protein antigen formulated in AddaVax to that elicited by the same antigen mixed with empty LNPs or by an antigen-matched mRNA-LNP vaccine, our data revealed that the sensing of the LNP component was the major driver for the transcriptional changes observed across immunization conditions. The sensing of the mRNA led to more limited transcriptional differences. An in-depth analysis of the LNP-driven transcriptional changes combined with the LIPSTIC system to track the antigen-presenting DCs highlighted selected clusters of cDC2s, whose transcriptomic profile was dictated by the sensing of the LNP rather than by their ability to engage T cells through CD40. Indeed, two biotin^lo^ clusters associated with LNP-containing immunization stood out for their heightened expression of costimulatory molecules (CD80 and CD86). This outcome is substantially different from what was seen using the LIPSTIC approach to label DCs in a tumor model, where the activation was restricted to the biotin^hi^ DCs that presented tumor antigens to CD4 T cells^37^. Taken together, these findings suggest that in the presence of a strong adjuvant, the need for T cell help to induce DC maturation is bypassed. LNP were also associated with genes encoding for molecules important for DC migration, including *Cd81*^41^, *Cd274*^42^, *Cxcr5*^43^, and *Gpr183*^45,65–67^. The last two were of particular interest for Tfh cell differentiation. Cxcr5 is a chemokine receptor that, in response to Cxcl13, mediates the localization toward and within B cell follicles^66^ and has been shown to play an instrumental role in DC localization and interaction with Cxcr5^+^ CD4 T cells at T-B borders^43^. Ebi2 (the protein encoded by *Gpr183*), allows cells to follow a 7α, 25-HC gradient that is enriched at T-B borders. Herein, we confirmed that LNP^+^ DCs, specifically cDC2s, have increased Cxcr5 expression, suggesting that intracellular sensing of LNP is involved in Cxcr5 upregulation. Furthermore, we demonstrated that the LNP component promotes an Ebi2/7α, 25-HC axis, by eliciting not only *Gpr183* expression by cDC2s but also the conversion of cholesterol into 7α, 25-HC. At the steady state, Ebi2 dictates the positioning of CD4 T cells at the periphery of the LN paracortex ^46^. It is believed that this spatial distribution facilitates antigen presentation by cDC2s, at least for antigens entering through the lymphatics, whose abundance exponentially decays with distance from the subcapsular sinus ^68^. It is important to keep in mind however that during immune responses Ebi2 is further upregulated by activated CD4 T cells, favoring their localization farther in the periphery of T cell zones, at T-B borders^17,46^. Splenic cDC2s also upregulate Ebi2 upon activation to relocate at the T-B borders^45^, suggesting that in LNP-based vaccination, cDC2s that have sensed LNP can better localize at T-B borders, where they can provide pro-Tfh signals, including soluble CD25, to antigen-specific CD4 T cells and guide their fate commitment to Tfh cells. In line with this model, loss of Ebi2 in CD4 T cells can have a profound disruption in Tfh cell responses^17^, and in LNP-based vaccination, we found a profound decrease in Tfh cell generation in C*h25h*^−/−^ mice that cannot produce the Ebi2 ligand 7α, 25-HC.

Our tracking studies also unveil that most cDC2s in the dLNs internalized mRNA-LNP vaccines by 24 hours after intramuscular administration. These data, combined with our photoactivation experiments demonstrating that the majority of the DCs internalizing mRNA-LNP were already in the dLN at the time of immunization, strongly suggest that mRNA-LNP can easily traffic through the lymphatics and are broadly processed by DCs in the dLNs. The rapid infiltration of innate cells in the muscle after the injection with nanoparticle-based vaccines combined with the local inflammation at the injection site supported the idea that vaccine uptake by infiltrating immune cells and migration of the vaccine-loaded cells to the dLN is critical in the initiation of adaptive immune responses^53,54^. This idea was further supported by the evidence that certain adjuvants such as Alum can persist at the injection site for several weeks^69^, slowly reach the dLN (~day 8) and are more efficiently internalized by DC in dLN^70^ after 7 days. This is in stark contrast to the rapid uptake of mRNA-LNP vaccines by the majority of DCs in the dLN at 24 hours shown by our work, which supports a model where the transportation of mRNA-LNP by DC from the injection site to the dLN makes only a small contribution to the total pool of mRNA-LNP found in the dLN within a day post-immunization.

Overall, in our study, we have uncovered a dual mechanism of adjuvanticity driven by mRNA vaccines and generated mechanistic knowledge that will pave the way to a targeted manipulation of the mRNA and LNP vaccine components for more effective mRNA-LNP vaccines.

### Limitations of the study

The ionizable lipid of the LNP component used in our study differs from those used in the licensed SARS-CoV-2 mRNA vaccines. Moderna utilizes the ionizable lipid SM-102 in the Spikevax® vaccine, and Pfizer the ionizable lipid ALC-0315 in the Comirnaty® vaccine, which are both different from the ionizable lipid used here^75^. It has been demonstrated by other groups that the difference in these ionizable lipids leads to downstream changes in tissue delivery and antibody responses, however, it remains unknown whether the mechanisms of Tfh cell induction are affected by the specific ionizable lipid used in vaccine preparations^71^. Future studies will be needed to determine if, and how, the ionizable lipids and other components of LNP impact the magnitude and quality of Tfh cell generation.

## ACKNOWLEDGEMENTS

M.L. was supported by the National Institute of Allergy and Infectious Diseases (NIAID; R01AI153064 and R01AI168312). E.B. was supported by the National Institute of Health (NIH; T32AI055400). N.P. was supported by the National Institute of Allergy and Infectious Diseases (NIAID, R01AI146101, R01AI153064, and P01AI158571). Z.L. was supported by the National Laboratory for Biotechnology (2022-2.1.1-NL-2022-00008) and the Hungarian Academy of Sciences (Lendület Program Grant (LP2017-7/2017)). G.D.V. was supported by NIH grants DP1AI144248 (Pioneer award) and R01AI173086. S.N.-H. was supported by a Bulgari Women & Science Fellowship and A.C. by a Damon Runyon Postdoctoral Fellowship. I.M. was supported by the NIAID (R01-AI091627), the National Cancer Institute (R01-CA278976), and a Commonwealth of Pennsylvania Health Research Formula Fund. S.C. was supported by the American Association of Immunologists Careers in Immunology Fellowship Program and the Charles A. King Trust Postdoctoral Research. A.R was supported by the National Institute of Allergy and Infectious Diseases (NIAID; 5R01AI158832 and R01AI155727).

We would like to thank Dr. Florin Tuluc and Jennifer Murray at the Children’s Hospital of Pennsylvania (CHOP) Flow Cytometry Core for technical assistance. R.S. would like to thank the UCLA QCB Collaboratory community directed by Matteo Pellegrini. Figures were created with the use of BioRender.com. We acknowledge the use of Grammarly, Inc for removing grammatical mistakes, spelling errors, and typos.

## AUTHOR CONTRIBUTIONS

E.B. and M.L. conceptualized the study and wrote the manuscript, with help from co-authors. E.B. and M.L. analyzed and graphed data. E.B. and G.P. produced figures. H.M. and N.P. produced mRNA vaccines. Y.T. produced LNP and mRNA-LNP. E.A. and Z.L. produced recombinant proteins. S.N-H. and G.D.V. conducted experiments with LIPSTIC mice. A.C. and G.D.V. conducted photoactivation experiments. J.C. performed human PBMC *in vitro* studies. E.B. and D.C. performed ICS assays. G.P. and R.S. performed a bioinformatic analysis of transcriptomic data. T.M. performed HAI assays. I.M. and G.D.V. provided feedback on the study design and data interpretation. S.C. and A.R. performed transwell assays and experiments with *Ch25h^−/−^* mice. E.B. performed all other experiments and analyses. M.L. supervised the study.

## STAR METHODS

### CONTACT FOR REAGENT AND RESOURCE SHARING

Further information and requests for resources and reagents should be directed to and will be fulfilled by the Lead Contact, Michela Locci.

### EXPERIMENTAL MODEL AND SUBJECT DETAILS

#### Mice

C57BL6/J, Cd11c-Cre-GFP (C57BL/6J-Tg(Itgax-cre,-EGFP)4097Ach/J), IFNAR^flox/flox^ (B6(Cg)-Ifnar1^tm1.1Ees^/J) were purchased from The Jackson Laboratory. *Cd40^G5^* and *Cd40lg^SrtA^*mice were generated as previously described^36^. *Ch25h^−/−^* were purchased from The Jackson Laboratory and maintained as *Ch25h^-/+^* x *Ch25h^-/+^* breeding. *Ch25h^−/−^* and *Ch25h^+/+^* littermates control were used. CAG-KikGR-transgenic mice^56^ were a gift from A. Hadjantonakis (Memorial Sloan Kettering Cancer Center). CAG-KikGR transgenic mice were back-crossed to the C57BL6 background for at least 10 generations at Rockefeller University. In all experiments, mice were immunized between 6 – 10 weeks of age.

Mice were housed in an Association for Accreditation of Laboratory Animal Care (AAALAC)-accredited facility and studies were conducted under protocols approved by the University of Pennsylvania’s Institutions Animal Care and Use Committee (IACUC).

### MATERIALS AND METHODS

#### Production of nucleoside-modified mRNA

Nucleoside-modified mRNA was produced as described previously^8^. Briefly, the coding sequences of the hemagglutinin (HA) of A/Puerto Rico/8/1934 (PR8), the receptor binding domain (RBD, amino acids 1-14 fused with amino acids 319-541) of SARS-CoV-2 (Wuhan-Hu-1, GenBank: MN908947.3), RBD fused via a SGGGG linker to the OTII binding region of ovalbumin (RBD-OVA, amino acids 318-340), extracellular domain of CD90.1 (Thy1.1, GenBank: AAR17087.1), or enhanced green fluorescent protein (eGFP) were codon-optimized, synthesized and cloned into the mRNA production plasmid as previously described^72^.

#### Lipid nanoparticle formulation of mRNA

Purified mRNAs were encapsulated in lipid nanoparticles using a self-assembling ethanolic lipid mixture of an ionizable cationic lipid, 1,2-distearoyl-*sn*-glycero-3-phosphocholine, cholesterol, and a polyethylene glycol-lipid as previously described^8^. This mixture was rapidly combined with an aqueous solution containing mRNA at acidic pH. The ionizable cationic lipid (pKa in the range of 6.0-6.5), proprietary to Acuitas Therapeutics (Vancouver, Canada) and LNP composition are described in the patent application WO 2017/004143. The average hydrodynamic diameter was ~80 nm with a polydispersity index of 0.02-0.06 as measured by dynamic light scattering using a Zetasizer Nano ZS (Malvern Instruments Ltd, Malvern, UK) and an encapsulation efficiency of ~95% as determined using a Quant-iT Ribogreen assay (Life Technologies). DiI-labled LNP were prepared by incorporating 1% of 1,1′-dioctadecyl-3,3,3′,3′-tetramethylindocarbocyanine perchlorate (DiIC18). Empty LNP used as adjuvants with recombinant protein and LNP encapsulating mRNA was performed by Acuitas Therapeutics (Vancouver, Canada) as described^72^.

#### Production of recombinant proteins

##### RBD and RBD-OVA

DNA sequences encoding the signal peptide (aa 1-14) and the RBD (aa 319-541) of SARS-CoV-2-Spike surface glycoprotein (NCBI Reference Sequence: YP_009724390) in fusion with a C-terminal hexahistidine-tag and a stop codon (hereafter referred to as RBD), or in fusion with an SGGGG linker with the OTII binding region of the chicken ovalbumin precursor protein (aa 318-340; NCBI Reference Sequence: NP_990483.2) followed by a C-terminal hexahistidine-tag and a stop codon (hereafter referred to as RBD-OVA) were optimized to mammalian codon preference, produced by gene synthesis (GenScript), and sub-cloned into the pCDNA3.1(-) mammalian expression plasmid. Recombinant RBD and RBD-OVA were produced in Expi293F mammalian cells, respectively, and affinity-purified from cell culture supernatants following the protocol described in detail in a prior study^73^. Purified proteins were buffer exchanged to phosphate buffered saline (PBS) and concentrated to 1 mg/ml, filter sterilized, and flash frozen in liquid nitrogen. The integrity and purity of proteins were tested by reducing and non-reducing SDS-PAGE and western blotting techniques.

##### rHA

Soluble and homotrimeric recombinant hemagglutinin (hereafter referred to as rHA) was synthesized in insect cells using a modified protocol previously detailed by^74^. The DNA sequence encompassing the signal peptide (amino acids 1-17) and the ectodomain (amino acids 18-529) of the hemagglutinin of Influenza A virus (A/Puerto Rico/8/1934 (PR8), GenBank No: ADX99484.1) in fusion with the thrombin cleavage site (RS*LVPRGSP*), followed by the foldon trimerization domain of the T4 bacteriophage fibritin (*GSGYIPEAPRDGQAYVRKDGEWVLLSTFL*, according to Stevens *et al*.^75^ and a C-terminal hexahistidine-tag and stop codon was produced by gene synthesis (GenScript), and sub-cloned into the pFastBac-HTA vector. Bacmids were created in DH10Bac *E. coli* cells following the guidelines of the Bac-to-Bac Baculovirus Expression System manual. Recombinant baculovirus was then generated and produced in *Spodoptera frugiperda* Sf9 insect cells using ExpiFectamin Sf transfection reagent as per the manufacturer’s instructions. The cells were cultured at 27°C in Sf-900 III serum-free medium in 6-well plates for 96 hours. The resulting cell culture supernatant, containing the P1 baculovirus rHA stock, was collected by centrifugation (2000 *xg*, 10 min 4°C). Subsequently, 0.5 ml P1 stock was employed to infect Sf9 cells in 6-well plates with approximately 60-70 % confluency. The cell culture supernatant (P2 baculovirus rHA stock) was collected 72 hours post-infection via centrifugation (2000 *xg*, 10 min 4°C). For the next passage, 0.5 ml of P2 rHA baculovirus was used to infect Sf9 cells in T75 flasks with a similar confluency. After 72 hours post-infection, the cell culture supernatant (P3 working baculovirus rHA stock) was collected, centrifuged (2000 *xg*, 10 min, 4°C), and filtered. The recombinant HA was expressed in *Trichoplusia ni* HighFive insect cells cultured in HyClone SFX-Insect medium as follows: HighFive cells from six confluent T175 flasks were harvested, collected by centrifugation (1200 x g, 24°C, 10 min), and then gently mixed with 15 ml of P3 baculovirus HA working stock. The mixture was incubated at room temperature for 20 minutes and slowly added to 210 ml of HyClone SFX-Insect medium in 1L PETG tissue culture Erlenmeyer flasks (Nalgene, #4115-1000). The cells were grown at 28°C on a shaker at 75 r.p.m. for 72 hours. The cell culture supernatant, containing the secreted homotrimeric rHA protein, was collected by centrifugation (2000 *xg*, 4°C, 10 min), filtered, and then subjected to affinity purification of HA following the steps outlined in the above-detailed RBD-purification protocol^76^. The purified rHA was buffer exchanged to PBS, concentrated to 1 mg/ml, filter-sterilized, and flash-frozen in liquid nitrogen. The integrity and purity of the homotrimeric rHA were assessed using reducing and non-reducing SDS-PAGE and western blotting techniques.

#### Immunizations and Injections

##### Immunizations

All immunizations were administered intramuscularly in the gastrocnemius muscle using a 0.5 mL 28 G x 1/2’’ insulin syringe (BD Biosciences, 329461). Unless otherwise stated, mRNA-LNP and recombinant proteins were administered at doses of 30 μg of antigen (mRNA or protein) and LNP were administered as the volume equivalent to 30 μg of mRNA-LNP (typically equivalent to 30 μl at 30 μg/μl). Prior to immunizations, mRNA-LNP, LNP, and/or recombinant proteins were diluted in PBS to a final volume of 50 μL.

##### Antibody Treatments

For IFNAR blocking experiments, 500 μg of anti-IFNAR blocking antibody was delivered intraperitoneally 24 hours prior to immunization.

#### Sample Collection and Processing

##### LN Processing

For DC-related analysis, draining LNs (inguinal and popliteal) were harvested in DMEM +10% FBS +1X Glutamax +1X Penicillin-Streptomycin, disrupted with scissors, and then digested in RPMI + 2% FBS + 20mM HEPES + 400U/mL type-IV collagenase for 30 minutes at 37°C. LNs were then passed through a 21G x 1 syringe (BD 309624) 5 times before being filtered through a 40uM mesh filter. For all other analyses, LNs were harvested and homogenized with a syringe plunger and filtered through a 40uM cell strainer on ice to create a single-cell suspension. *Muscle Processing.* The gastrocnemius muscle was harvested DMEM +10% FBS +1X Glutamax +1X Penicillin-Streptomycin, disrupted with scissors, and then digested in RPMI + 7000U Collagenase II and 10% FBS for 1.5 hours at 37°C. Samples were then pelleted at 400xg for 5 minutes and the supernatant was discarded. Pellets were resuspended in RPMI + 500U Collagenase II and 50U Dispase for 30 minutes at 37°C. Samples were then passed through a 21G x 1 syringe (BD 309624) 5 times before being filtered through a 40uM mesh filter to create a single-cell suspension.

##### Spleen Processing

Spleens were collected, homogenized with a syringe plunger, and filtered through a 70uM filter to generate a single-cell suspension. Red blood cells were lysed with ACK lysis buffer for 5-8 minutes on ice and the reaction was stopped with cold PBS. Cells were resuspended in PBS supplemented with 2% FBS + 1mM EDTA.

##### Bone Marrow (BM) Processing

BM was flushed from both femurs and tibias from each mouse using a 1 mL 25 G x 5/8’’ syringe. RBCs were lysed as described above.

##### Serum Processing

Following collection, blood was centrifuged at 14,000 x g (maximum speed) for 30 minutes, 4°C. Serum was recovered and stored at −20°C for ELISAs.

All organs were kept on ice and immediately processed after collection.

#### T cell intracellular cytokine assay

This assay was performed as previously described^9^. Briefly, upon sample processing, CD4 T cells were isolated from draining lymph nodes using the Mojosort^TM^ Mouse CD4 T cell isolation kit according to the manufacturer’s instructions. 10^6^ CD4 T cells were incubated in IMDM (+10% FBS, +1% Penicillin/streptomycin +2mM L-glutamine +1mM sodium pyruvate + 55 uM 2-mercaptoethanol) containing PMA (50 ng/mL) and ionomycin (1 μg/mL). After 2 hours, GolgiPlug was added at 1:1000 and the cells were incubated for 3 additional hours. Unstimulated cells were incubated with GolgiPlug and without PMA and Ionomycin. After stimulation, cells were stained as described in the Flow Cytometry section.

#### Flow Cytometry

All staining steps were performed at 4°C in PBS + 2% FBS + 5mM EDTA. Prior to staining, single-cell suspensions were incubated with anti-mouse CD16/CD32 blocking antibody at 1:1000 (~7.6 μg/mL) for 20 minutes.

##### Tfh Cells

Cells were incubated with biotinylated anti-Cxcr5 for 1 hour. Cells were washed and then incubated with a cocktail of fluorescently labeled anti-mouse mAbs, streptavidin and Fixable Viability Dye e780 (**Table S1**) for 30 minutes. Cells were washed and then fixed and permeabilized in FoxP3/Transcription Factor staining buffer set, according to the manufacturer’s instructions, before intranuclear staining with anti-BCL6.

##### GC B Cells

Cells were incubated with biotinylated anti-CD138 for 1 hour. Cells were washed and then incubated with a cocktail of fluorescently labeled anti-mouse mAbs, streptavidin and Fixable Viability Dye e780 (**Table S2**) for 30 minutes. Samples were then washed before fixation with 1% paraformaldehyde for 30 minutes.

##### APCs

Cells were incubated with biotinylated anti-Cxcr5 for 1 hour. Cells were washed and then incubated with a cocktail of fluorescently labeled anti-mouse mAbs, streptavidin and Fixable Viability Dye e780 (**Table S3**) for 30 minutes.

##### DCs, MBCs, and Human PBMCs

Cells were washed and then incubated for 20-30 minutes with a cocktail of viability dye and the corresponding fluorescently labeled antibody cocktail in FACS buffer (PBS containing 2% FBS and 1mM EDTA): *DCs* (**Tables S4 and S5**)), *MBCs* (**Table S6**), *Human PBMCs* (**Table S7**).

##### Intracellular staining for cytokines

Upon Fc blockade, cells were fixed and permeabilized with Cytofix/Cytoperm according to the manufacturer’s instructions. Cells were then resuspended with IL-21-Fc chimera protein (1:20) for 30 minutes, washed, and resuspended in anti-human IgG PE mAb for 20 minutes. Next, cells were washed and incubated overnight at 4°C in a mixture of the remaining antibodies (**Table S8**). The following day the cells were washed prior to acquisition.

##### LIPSTC

Lymphocytes from draining lymph nodes were incubated for 30 min at 4°C temperature with a cocktail of surface marker antibodies (**Table S9**) and Live, B220^−^CD3^−^NK1.1^−^ MHCII^hi^ CD11c^+^ DCs were sorted and used for single-cell transcriptomic studies.

All flow cytometry samples were acquired on a Cytek Aurora using SpectroFlow v2.2 (Cytek) or an LSR Fortessa (BD). Data were analyzed using FlowJo v.10.7.2 (Treestar).

#### LIPSTIC Labeling

Upon sample processing, splenocytes from *CD40lg*^SrtA/Y^CD4-Cre OT-II mice were incubated using a cocktail of biotinylated antibodies targeting Ter119, CD11c, CD25, B220, NK1.1 and CD8, followed by negative selection using anti-biotin beads (Miltenyi), as per the manufacturer’s instructions. 3 × 10^5^ *CD40lg*^SrtA/Y^CD4-Cre OT-II CD4^+^ T cells were transferred into *Cd40*^G5/G5^ recipient mice 18-hours prior to immunization. 22-hours post-immunization, Biotin-LPETG was injected intraperitoneally (i.p.) (100 μl of 20 mM solution in PBS) six times 20 minutes apart, and then inguinal and popliteal lymph nodes were harvested 20 minutes after the last injection. Single-cell suspensions were incubated with 1 μg/ml anti-CD16/32 (2.4G2, BioXCell) in PBS supplemented with 0.5% BSA and 2 mM EDTA (PBE) for 5 minutes at room temperature and then stained for cell surface markers at 4 °C for 20 minutes in PBS. For single-cell transcriptomic analysis, stained cells were further incubated with DNA-barcoded anti-biotin and sample hashtag (anti-MHC-I) antibodies (BioLegend) for 20 minutes in PBE, washed three times with PBE, and bulk-sorted.

#### Single-cell RNA sequencing (scRNA-seq) library preparation

Sorted DCs were collected into a microfuge tube with 300 μl PBS supplemented with 0.4% BSA. After the sort, tubes were topped with PBS 0.4% BSA, centrifuged, and the buffer was carefully reduced by removing the volume with a pipette to a final volume of 40 μl. Cells were counted for viability and immediately submitted to library preparation. The scRNA-seq library was prepared using the 10X Single Cell Chromium system, according to the manufacturer’s instructions, at the Genomics Core of Rockefeller University and was sequenced on an Illumina NovaSeq SP flowcell to a minimum sequencing depth of 30,000 reads per cell using read lengths of 26 bp read 1, 8 bp i7 index, 98 bp read 2.

#### Analysis of scRNA-seq data

ScRNA-seq reads were processed with CellRanger v6.1.2. A custom reference genome was created by adding the RBD mRNA sequence to the GRCm38 mouse genome. Individual count tables were read into R and downstream analysis was performed using Seurat v4^77^. Sample demultiplexing was performed using the HTODemux function and barcodes identified as negative or doublets were removed. Cells with at least 500 genes and <5% mitochondrial counts were considered for analysis, resulting in a total of 17,015 genes and 12,788 cells across the three treatments. The PCA plot was generated using DESeq2 from pseudo-bulk counts aggregated by individual animals. The biotin signal was normalized using the NormalizeData function (normalization.method = “CLR”, margin=1). Cells were classified as biotin-positive or -negative by using an elbow plot for ranking, where cells above the elbow (threshold set at 2.8) were considered positive and below were considered negative. GSEA was performed by contrasting pseudo-bulk counts of biotin positive and negative cells with DESeq2 and testing genes ranked by −log10 p-value (signed by log fold change) for enrichment of a previously published LIPSTIC signature^37^ with the R package fgsea. Single-cell transcriptomic counts were normalized using sctransform, while regressing out UMI counts and the percentage of mitochondrial counts.

Canonical correlation analysis (CCA) as implemented by Seurat v4 was used for treatment integration. Integrated data underwent principal component analysis (PCA) and graph-based clustering was then performed using the FindNeighbors and FindClusters functions, resulting in 13 clusters. Three clusters were identified as contaminants based on the expression of canonical marker genes: macrophages (Mafb), T cells (Cd3d) and B cells (Cd79b). These clusters were removed, and the remaining cells (n=12,031) were re-clustered following the same workflow just described, yielding 14 clusters. Cluster markers were identified using the FindAllMarkers function. DC annotation was performed based on a curated list of marker genes, as shown in Figure 3E.

ISG (genes: *Ifit3b, Ifi213, Ifit2, Tnfsf10, Cmpk2, Cxcl10, Rsad2, Cd69, Gbp6, Ifit1*) scores were computed using the AddModuleScore function of Seurat. Tfh (genes: *Cd80, Cd86, Cd274, Il2ra, Gpr183*) and LIPSTIC (gene set from^37^) scores were computed using UCell^78^, with kNN smoothing.

#### Human *in vitro* experiments

Cryopreserved PBMC samples were thawed and washed in complete RPMI (RPMI, 10% FBS, 1% Pen Strep, 1% GlutaMAX). 2×10^6^ cells/well were plated in a 48-well plate. eGFP-mRNA DiI-LNP were added at a final concentration of 10 μg/mL and incubated at 37 °C, 5% CO_2_ for the indicated time periods before flow cytometry staining and data acquisition.

#### KikGR Photoconversion

Mice were anesthetized with isoflurane, shaved, and washed with ethanol and 0.005M iodine solution. Immediately prior to immunization, photoactivation was performed by exposing either the gastrocnemius muscle or inguinal LN to 415nM light for two minutes, unless otherwise indicated. The incision was then closed using FST autoclips.

#### Transwell Assay

This assay was performed as previously described^49^. Briefly, lipids from LNs were extracted using the Folch method for lipid extraction^79^. Lyophilized lipid was dissolved at 100 mg ml−1 in ethanol. Lipid extracts were then diluted in 10 volumes of sterile chemotaxis medium (RPMI + 0.5% fatty acid-free BSA) and tested for GPR183-dependent bioactivity by seeding on transwell, 50:50 mixed, M12 B cell line transduced with an GPR183–IRES–GFP retroviral construct and mock M12 cells. The migration assay was performed at 37 °C for 3 h and cells were analyzed by flow cytometry. The migration of GPR183–GFP+ M12 cells over M12 cells (which indicate the relative concentration of the GPR183 ligand, 7α,25-HC) was normalized to the migration toward lipid-free migration medium and indicated in the text as ‘relative migration (a.u.)’. The purified 7α,25-HC was used as a positive control at a concentration of 100 nM.

#### qPCR

Whole LNs were extracted into RTL buffer and lysed using the Qiagen TissueLyzer® II system. Tissues were disrupted using Qiagen 5 mm steel beads under the following conditions: 2X 30 Hz, 30 s per cycle. Total RNA was extracted using a Qiagen RNeasy kit following the manufacturer’s instructions. cDNA was then transcribed using Superscript II reverse transcriptase according to the manufacturer’s instructions. Quantitative real-time PCRs (qPCRs) for *Gapdh*, *Ch25h*, *Cyp7b1*, and *Hsd3b7* were performed using the following primer pairs: *Gapdh*: forward 5’GCACAGTCAAGGCCGAGAAT-3’ and reverse 5’-GCCTTCTCCATGGTGGTGAA-3’^80^; *Ch25h*: forward 5’-GCGACGCTACAAGATCCA-3’ and reverse 5’-CACGAACACCAGGTG CTG-3’; *Cyp7b*: forward 5’-TTCCTCCACTCATACACAATG-3’ and reverse 5’-CGTGCTTTTCTTCTTACCATC-3’; *Hsd3b7*: forward 5’-ACCATCCACAAAGTCAACG-3’ and reverse 5’-TCTTCATTGCCCCTGTAGA-3’^48^. qPCRs were set up using SYBR Green PowerUp Master Mix and ran on QuantStudio 6 PCR System. Relative mRNA levels were calculated by 2^-ΔCt^ according to *Gapdh* gene abundance.

#### ELISPOT

MultiScreen HTS IP filter plates of 0.45 mm were coated with 2.5 mg/mL recombinant HA in bicarbonate buffer (35 mM NaHCO_3_ and15 mM Na_2_CO_3_) overnight (ON) at 4°C. Plates were washed 3X with PBS and blocked with complete DMEM (10% FBS + 1X Glutamax) for 2 hours at room temperature (RT). Single-cell suspensions of BM cells were serially diluted in complete DMEM with halving dilutions starting at one million cells. Following ON incubation at 37°C, 5% CO_2_ plates were washed 3X with wash buffer (0.05% Tween-20 in PBS). HRP-conjugated IgG was diluted 1:1000, IgG1 and IgG2c were diluted 1:3,000 in complete DMEM for 2 hours at RT. Plates were washed 3X with wash buffer and spots were developed using BD ELISPOT AEC Substrate Set. Membranes were dried ON and counted using a CTLImmunospot analyzer.

#### ELISA

Nunc Maxisorp flat-bottom 96 well plates were coated with 1 mg/mL recombinant HA in bicarbonate buffer overnight (ON) at 4°C. Plates were washed 3X with wash buffer and blocked with blocking buffer (2% bovine serum albumin in PBS) for 1 hour at RT. Serum samples were serially diluted in blocking buffer and incubated for 2 hours at RT then washed 3X with wash buffer. HRP-conjugated IgG was diluted 1:1000, IgG1 and IgG2c were diluted 1:5,000 in blocking buffer and incubated for 1 hour, then washed 3X with wash buffer. Plates were developed with Pierce TMB Substrate and the reaction was stopped with 2 N sulfuric acid. Absorbance was measured at 450 nm using a Tecan Infinite 200Pro. Area under the curve (AUC) was calculated using Prism v10.1.1 with the total peak area being reported.

#### Hemagglutination Inhibition (HAI) Assay

This assay was performed as previously described^8^. Briefly, heat-inactivated sera were diluted 1:20 in PBS, then serially diluted 1:2 in 50 μL in 96-well U-bottom plates. Four hemagglutinating doses of A/Puerto Rico/8/1934 virus were added, followed by 12.5 μL of turkey erythrocyte solution (final volume 125μL). Samples were incubated for 45 minutes at RT prior to being scanned. HAI titers were determined as the highest dilution of the sample that inhibited four agglutinating doses of the influenza virus.

#### Statistical Analysis

GraphPad Prism v10.1.1 was used to conduct all statistical analysis. Shapiro-Wilk and Kolmogorov-Smirnov tests were performed to establish the normal distribution of the data. The statistical analyses used are described in each figure legend and include the following calculations of statistical differences according to the distribution of the data: Two-way ANOVA with Tukey’s correction for multiple comparisons; One-way ANOVA Kruskal-Wallis test with Dunn’s correction for multiple comparisons; Unpaired two-tailed Mann-Whitney *U* test, with or without a two-stage Benjamini, Krieger and Yekutieli FDR of 1% to correct for multiple comparisons; One-tailed paired Wilcoxon test. The precise number of samples analyzed in each graph is reported in the figure legends. Statistical significance was set at the critical values of p < 0.05 (*), p < 0.01 (**), p < 0.001 (***), and p < 0.0001 (****). Data were expressed as an average +/- standard error of the mean (SEM).

## SUPPLEMENTAL INFORMATION

**Supplementary Figure 1. IFNAR expression on DCs regulates the magnitude of the GC response to mRNA-LNP, related to Figure 1**.

**(A)** Tfh cell (Live, B220^−^CD4^+^CD44^+^CD62L^−^Cxcr5^+^Bcl6^+^) gating strategy.

**(B)** GC B cell (Live, CD19^+^CD3^−^FAS^+^GL7^+^) gating strategy.

**(C)** Control (CD11c-cre^+^) or IFNAR cKO (CD11c-cre IFNAR^flox/flox^) mice received a single IM immunization. IL-6 levels were determined in the dLN extracts 4 hours post-immunization.

In (A-B), mice received a single IM immunization with 30 μg of influenza virus hemagglutinin (HA) mRNA-LNP. In (C) mice received a single IM immunization with 30 μg of RBD mRNA-LNP. In (C), n = 5-6 mice per group. Data is compiled from 2 independent experiments. Statistical analysis: Unpaired two-tailed Mann-Whitney *U* test was performed. Error bars represent mean + SEM.

**Supplementary Figure 2. mRNA-LNP drive a Th1-biased immune response, related to Figure 2**.

**(A)** Gating strategy for IFN-γ, IL-4 and IL-21 production by total CD4 T cells and Tfh cells determined by flow cytometry.

**(B)** Gating strategy for Tfh cells. Tfh cells were gated based on Cxcr5 expression. Naïve (CD44^−^, black) CD4 T cells were used as a negative control to set the Tfh cell gate.

**(C and D)** Representative IL-21 expression (**C**) and quantification of IL-21^+^ Tfh cells (**D**) measured by flow cytometry.

**(E)** Gating strategy for MBCs (Live, CD19^+^Fas^−^IgD^−^IgM^−^CD27^−^CD38^+^). HA-specific MBCs were defined based on the co-expression of two HA probes (BV421- and AF488-conjugated).

In (**A-D**), mice received a single IM immunization with 30 μg of RBD mRNA-LNP or 30 μg of recombinant RBD combined with LNP, as indicated. In (**E**) mice received a single IM immunization with 30 μg of HA mRNA-LNP. In (**A**-**E**), n = 9-10 mice per group. Data were combined from 2-3 independent experiments. Statistical analysis: unpaired two-tailed Mann-Whitney *U* test was conducted. Error bars represent mean + SEM.

**Supplementary Figure 3. LIPSTIC gene signature is enriched in biotin^+^ DCs, related to Figure 3**.

**(A)** Amino acid sequence of the RBD-OVA construct, consisting of a signal peptide (black) to facilitate secretion (absent in the secreted protein), the RBD portion of SARS-CoV-2 Spike (red), a GS linker (blue), the peptide sequence of OVA recognized by OT-II (amino acids 318-340, green) and the hexahistidine-tag (purple) for affinity purification.

**(B)** Schematic of the RBD-OVA mRNA-LNP construct.

**(C)** UMAP clustering of sequencing data from Figure **3** before integration.

**(D)** Feature plot of biotin expression before integration, displayed as a continuous variable.

**(E)** Heatmap displaying the top 10 differentially regulated genes in each DC group/cluster.

**(F)** Elbow plot of biotin-labeling, where the elbow represents the threshold (2.8) for biotin^+^ cells.

**(G)** GSEA analysis showing the enrichment of LIPSTIC-signature genes in biotin^+^ DCs.

**(H and I)** *Cxrc5* expression in each treatment group (**H**) and on the different DC clusters (**I**).

Mice were immunized with either 30 μg of RBD-OVA mRNA-LNP, 30 μg recombinant RBD-OVA combined with LNP, or 30 μg recombinant RBD-OVA combined with AddaVax; n= 2 mice per group.

**Supplementary Figure 4. LNP drive a pro-Tfh cell signature in cDC2s, related to Figure 4**.

**(A)** Representative CD25 expression in cDC1s and cDC2s (as indicated) at baseline (0 hours, black) versus 24 hours post-immunization (blue), determined by flow cytometry. Fluorescent intensity is displayed as count (normalized to mode).

Mice received a single IM immunization containing 20 μg of fluorescent mRNA-LNP.

**Supplementary Figure 5. Conventional DCs are capable of mRNA-LNP uptake and translation, related to Figure 5**.

**(A)** Gating strategy for analyzing APC in mice. Live, TCRβ^−^CD19^−^ were stratified into LCs (CD11c^+^MHCII^+^EpCAM^+^), cDC1s (CD11c^+^MHCII^+^EpCAM^−^CD103^+^CD11b^−^), cDC2s (CD11c^+^MHCII^+^EpCAM^−^CD103^−^CD11b^+^), pDCs (CD11b^−^PDCA1^+^), inflammatory monocytes (CD11b^+^PDCA1^−^Ly6G^−^SiglecF^−^Ly6C^+^) and macrophages (CD11b^+^PDCA1^−^ Ly6G^−^SiglecF^−^Ly6C^−^).

**(B)** Quantification of LNP binding/uptake (DiI^+^) and mRNA translation (eGFP^+^) in the indicated cell populations, as defined in **A**. Each vertical bar represents 100% of the indicated cell type and the color represents the portion of cells that are DiI^+^ eGFP^+^ (green), DiI^+^ eGFP^−^ (red), and DiI^−^ eGFP^−^ (blue). Error bars represent SEM.

**(C)** Quantification of cell types from **A**, represented as a percentage of Live, Linage (Lin)^−^ cells at the indicated time points.

**(D)** Full gating strategy for mouse DCs, used to define DC populations in **Figure 5B-F**.

**(E)** Gating strategy for human DCs, used to define DC populations in **Figure 5G**.

**(F)** Quantification of uptake and translation of mRNA-LNP in human DCs from **Figure 5G**, displayed as individual donors.

For (**A-D**) mice were immunized with 20 μg of eGFP mRNA-LNP-DiI. For (B and C), n=6 mice per time point. Data is combined from two independent experiments. Error bars represent mean + SEM. *p ≤ 0.05, **p ≤ 0.01.

**Supplementary Figure 6. Photoactivation of muscle and lymph nodes reveals vaccine uptake within draining LNs, related to Figure 6**.

**(A)** Gating strategy for defining DCs after photoactivation (PA).

**(B)** Efficacy of photoactivation in the gastrocnemius muscle. Percentage of RFP^+^ DCs displayed as a percentage of total DCs (**Left**) or percentage of total CD45^+^ (**Right**) cells in the gastrocnemius immediately following photoactivation.

**(C)** Quantification of CD45^+^ cell (**Left**) and DC (**Right**) frequencies in the gastrocnemius muscle at the indicated time points post-vaccination.

**(D)** Quantification of data from **Figure 6D** and **E**, displaying RFP^+^ and RFP^−^ Thy1.1^+^ DCs as frequency of total DCs from the dLNs of PBS (control) and mRNA-LNP immunized mice.

In (**A and D-E**) mice were injected with 30 μg Thy1.1 mRNA-LNP. In (**C**) mice were injected with 20 μg of eGFP mRNA-LNP-DiI. In **(B)** n = 3-6 mice per group combined from 2 experiments. In **(C-E)** n = 5-6 mice per group combined from 2 experiments. Statistical analysis: In panels **(C and D-E)** a Two-way ANOVA with Tukey’s correction for multiple comparisons was performed. Error bars represent mean + SEM. *p ≤ 0.05, **p ≤ 0.01, ***p ≤ 0.001, ****p ≤ 0.0001.

