## Supplementary figures and images for "Distinct components of nucleoside-modified messenger RNA vaccines cooperate to instruct efficient germinal center responses"

### Supplementary Figure 1

#### Supplementary Figure 1

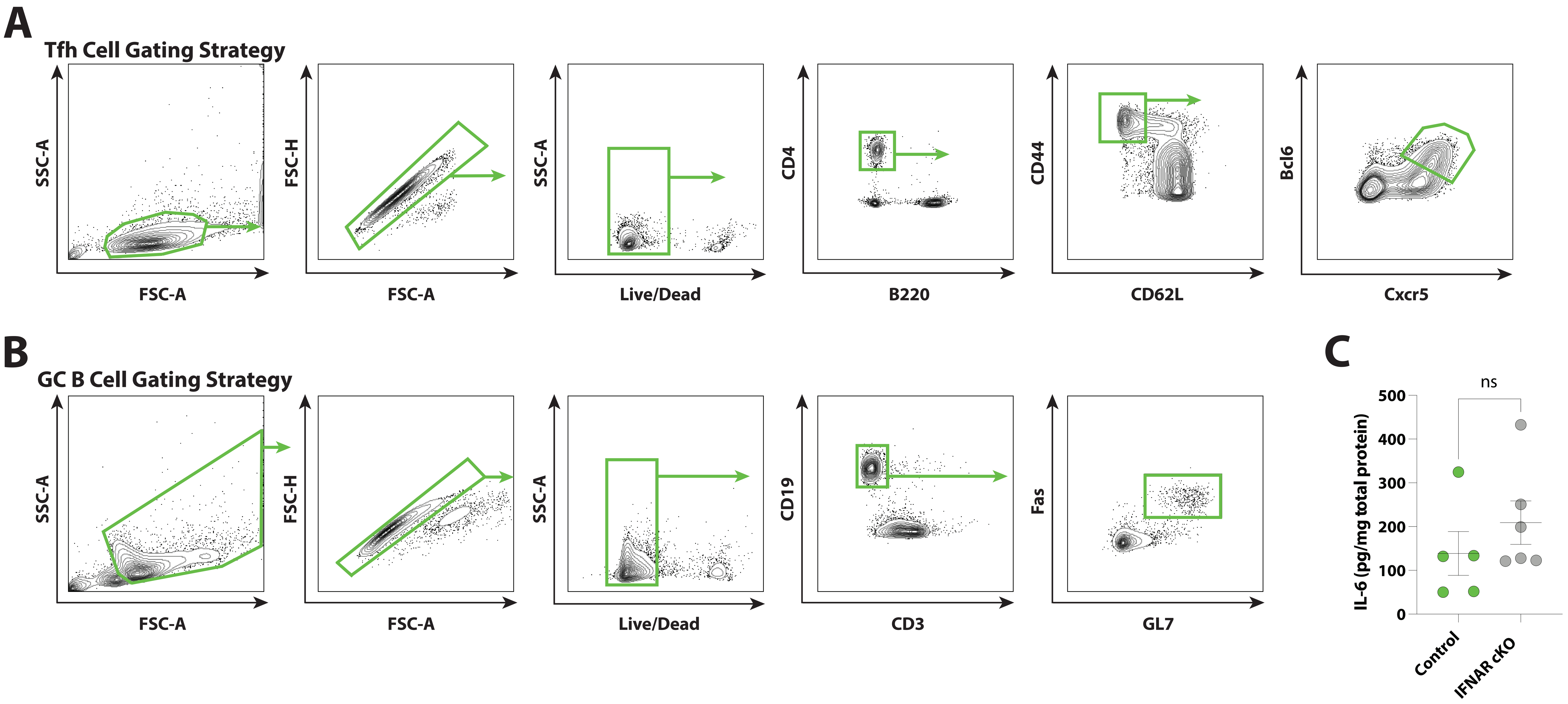

### Supplementary Figure 2

#### Supplementary Figure 2

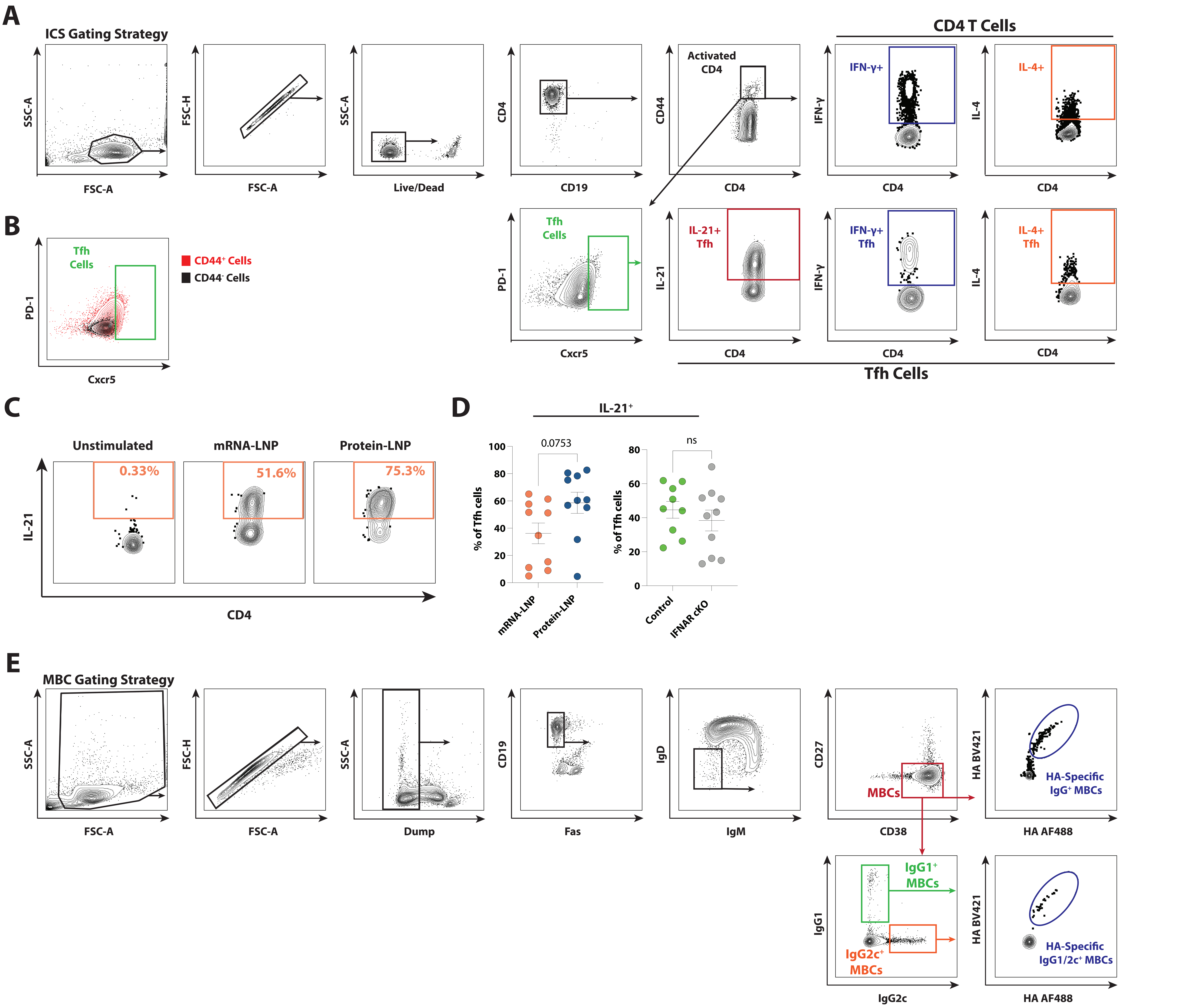

### Supplementary Figure 3

#### Supplementary Figure 3

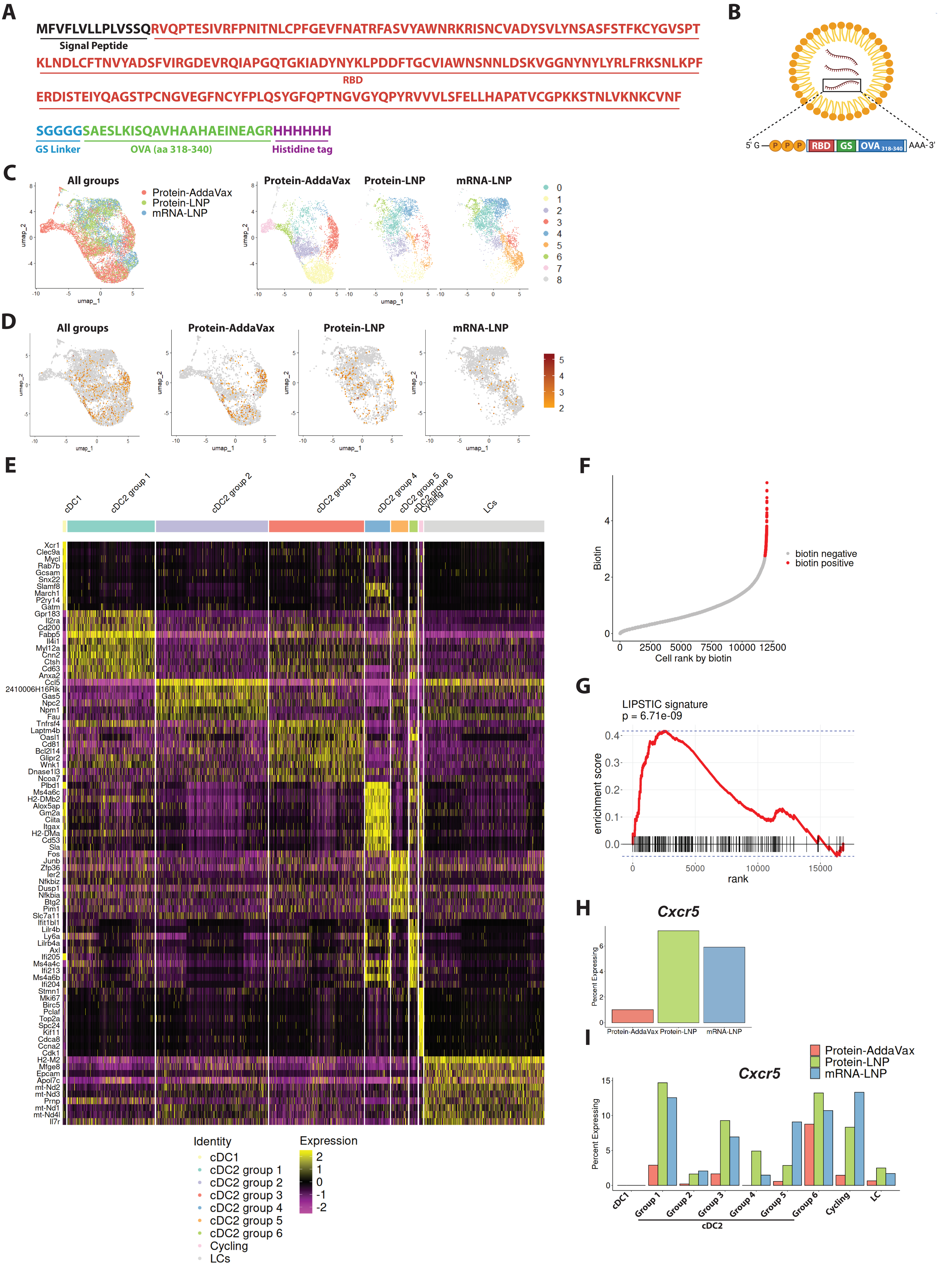

### Supplementary Figure 4

#### Supplementary Figure 4

**A**

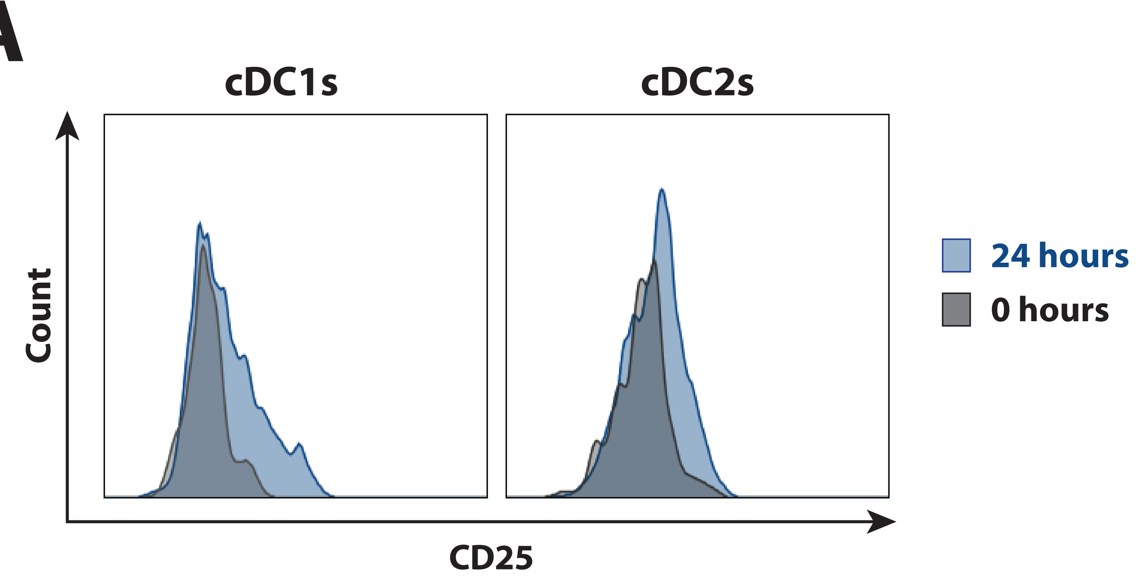

### Supplementary Figure 5

#### Supplementary Figure 5

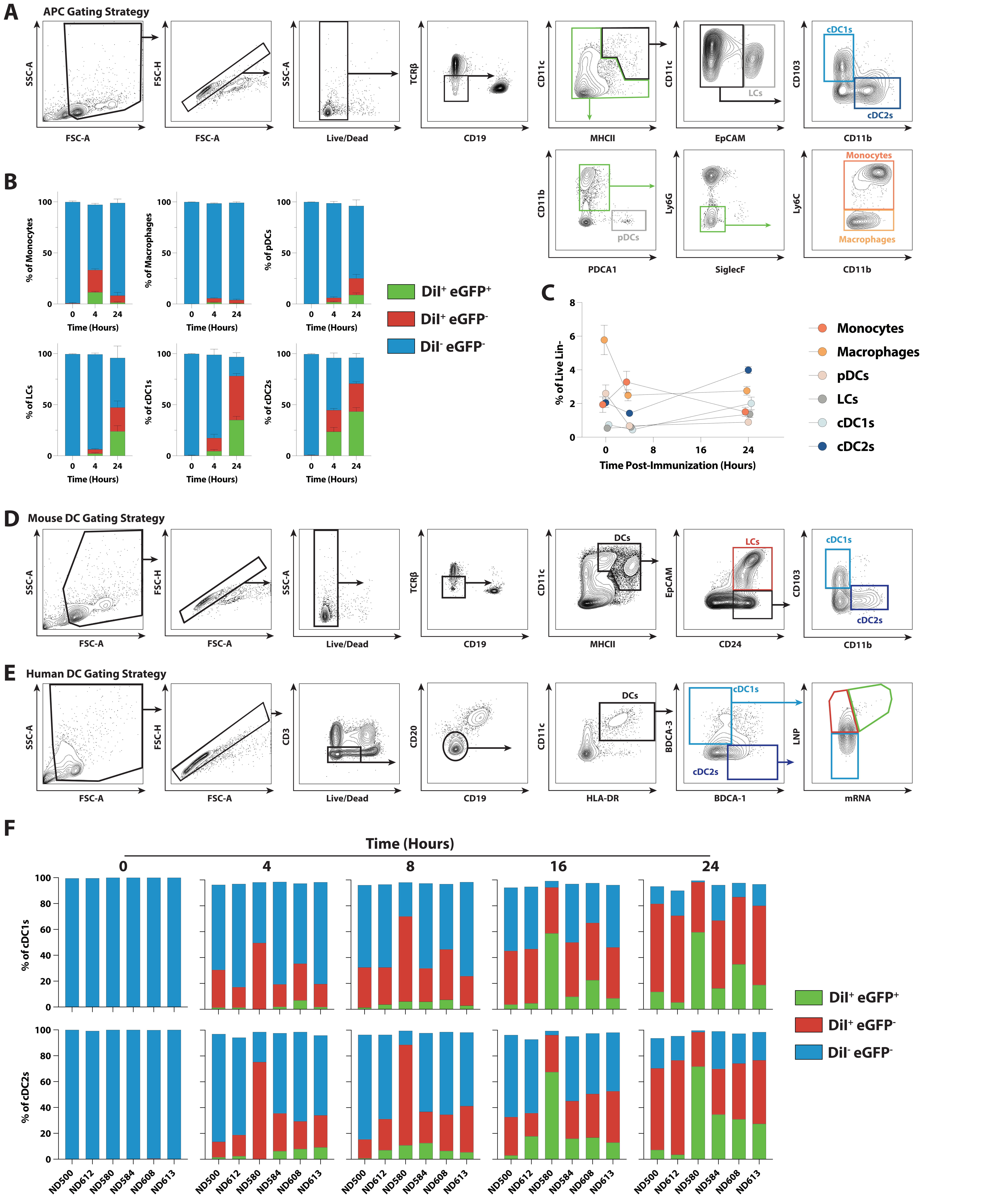

### Supplementary Figure 6

#### Supplementary Figure 6

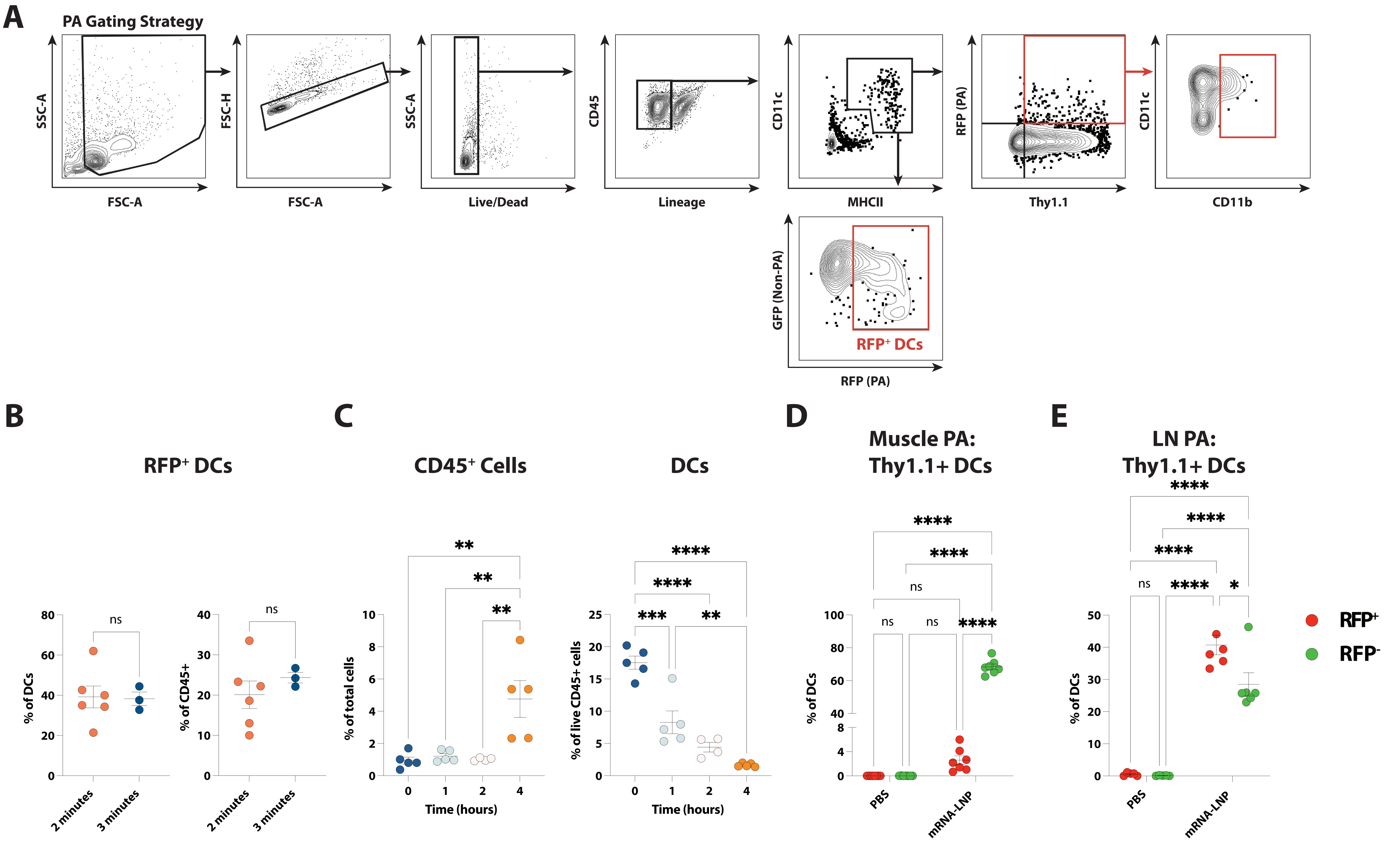
