## Supplementary Tables for "Distinct components of nucleoside-modified messenger RNA vaccines cooperate to instruct efficient germinal center responses"

| Stain | Fluorochrome | Clone | Vendor | Catalogue # | Dilution |
|---|---|---|---|---|---|
| CXCR5 | Biotin | SPRCL5 | eBioscience | 13-7185-82 | 1:50 |
| Streptavidin | BV421 | - | BioLegend | 405225 | 1:500 |
| B220 | BV650 | RA3-6B2 | BioLegend | 103241 | 1:400 |
| CD4 | PerCP-Cy5.5 | RM4-5 | BioLegend | 100540 | 1:200 |
| CD44 | BV605 | IM7 | BioLegend | 103047 | 1:400 |
| CD62L | BUV395 | MEL-14 | BD | 740218 | 1:400 |
| PD-1 | PE | RMP1-30 | BioLegend | 109104 | 1:200 |
| Bcl6 | AF647 | K112-91 | BD | 624024 | 1:200 |
| Live/Dead | eFluor 780 | - | eBioscience | 65-0865-14 | 1:2000 |

**Table S1. Tfh cell panel, related to Figures 1, S1, 4, 6.**

| Stain | Fluorochrome | Clone | Vendor | Catalogue # | Dilution |
|---|---|---|---|---|---|
| CD138 | Biotin | 281-2 | BD Biosciences | 553713 | 1:500 |
| Streptavidin | BV650 | - | BioLegend | 405232 | 1:200 |
| CD3e | BUV395 | 145-2c11 | BD Biosciences | 563565 | 1:100 |
| CD19 | BV605 | 6D5 | BioLegend | 115540 | 1:800 |
| GL7 | PerCP-Cy5.5 | GL7 | BioLegend | 144610 | 1:400 |
| FAS | BV510 | Jo2 | BD Biosciences | 563646 | 1:800 |
| IgD | PE-Cy7 | 11-26c.2a | BioLegend | 405720 | 1:400 |
| Live/Dead | eFluor 780 | - | eBioscience | 65-0865-14 | 1:2000 |

**Table S2. GC B cell Panel, related to Figures 1, S1, S4, 6.**

| Stain | Fluorochrome | Clone | Vendor | Catalog # | Dilution |
|---|---|---|---|---|---|
| CXCR5 | Biotin | SPRCL5 | eBioscience | 13-7185-82 | 1:50 |
| Streptavidin | BUV661 | - | BD | 612979 | 1:200 |
| MHCII | BUV805 | M5/114.15.2 | BD | 748844 | 1:1000 |
| CD11b | BV421 | M1/70 | BioLegend | 101236 | 1:800 |
| TCRb | BV510 | H57-597 | BioLegend | 109234 | 1:100 |
| PDCA1 | BV605 | 927 | BioLegend | 127025 | 1:100 |
| Ly6G | BV650 | 1A8 | BioLegend | 127641 | 1:200 |
| CD103 | BV785 | 2E7 | BioLegend | 121439 | 1:50 |
| Ly6C | PerCP-Cy5.5 | HK1.4 | Invitrogen | 45-5932-82 | 1:200 |
| CD19 | PE-Cy5 | 6D5 | BioLegend | 115510 | 1:800 |
| CD11c | PE-Cy7 | N418 | BioLegend | 117318 | 1:50 |
| SiglecF | APC | S17007L | BioLegend | 155508 | 1:200 |

| | | | | | |
|---|---|---|---|---|---|
| CD326 | AF700 | G8.8 | BioLegend | 118240 | 1:100 |
| Live/Dead | eFluor 780 | - | eBioscience | 65-0865-14 | 1:2000 |

**Table S3. APC panel, related to Figures S5A, B, and C.**

| Stain | Fluorochrome | Clone | Vendor | Catalog # | Dilution |
|---|---|---|---|---|---|
| MHCII | BUV805 | M5/114.15.2 | BD | 748844 | 1:1000 |
| CD86 | BV421 | GL-1 | BioLegend | 105032 | 1:200 |
| TCRb | BV510 | H57-597 | BioLegend | 109234 | 1:100 |
| CD24 | BV605 | M1/69 | BioLegend | 101827 | 1:400 |
| CD80 | BV650 | 16-10A1 | BioLegend | 104732 | 1:400 |
| CD103 | BV785 | 2E7 | BioLegend | 121439 | 1:50 |
| CD11b | FITC | M1/70 | BioLegend | 101205 | 1:800 |
| CD40 | PE | 3/23 | BioLegend | 124609 | 1:200 |
| CD19 | PE-Cy5 | 6D5 | BioLegend | 115510 | 1:800 |
| CD11c | PE-Cy7 | N418 | BioLegend | 117318 | 1:50 |
| CD326 | AF700 | G8.8 | BioLegend | 118240 | 1:100 |
| Live/Dead | eFluor 780 | - | eBioscience | 65-0865-14 | 1:2000 |

**Table S4. DC panel 1, related to figure 1G and H.**

| Stain | Fluorochrome | Clone | Vendor | Catalog # | Dilution |
|---|---|---|---|---|---|
| CD25 | BUV395 | PC61 | BD | 564022 | 1:400 |
| MHCII | BUV805 | M5/114.15.2 | BD | 748844 | 1:1000 |
| CD11b | BV421 | M1/70 | BioLegend | 101236 | 1:800 |
| TCRb | BV510 | H57-597 | BioLegend | 109234 | 1:100 |
| CD24 | BV605 | M1/69 | BioLegend | 101827 | 1:400 |
| CD80 | BV650 | 16-10A1 | BioLegend | 104732 | 1:400 |
| PD-L1 | BV711 | 10F.9G2 | BioLegend | 124319 | 1:800 |
| CD103 | BV785 | 2E7 | BioLegend | 121439 | 1:50 |
| CD8a | PerCP-Cy5.5 | 53-6.7 | BioLegend | 100734 | 1:200 |
| CD19 | PE-Cy5 | 6D5 | BioLegend | 115510 | 1:800 |
| CD11c | PE-Cy7 | N418 | BioLegend | 117318 | 1:50 |
| CD200 | AF647 | OX-90 | BioLegend | 123816 | 1:400 |
| CD326 | AF700 | G8.8 | BioLegend | 118240 | 1:100 |
| Live/Dead | eFluor 780 | - | eBioscience | 65-0865-14 | 1:2000 |

**Table S5. DC panel 2, related to Figures 5 and S5.**

| Stain | Fluorochrome | Clone | Vendor | Catalogue # | Dilution |
|---|---|---|---|---|---|
| IgG2b | BUV563 | R12-3 | BD Biosciences | 749138 | 1:200 |
| IgE | BUV805 | R35-72 | BD Biosciences | 749151 | 1:200 |

| | | | | | |
|---|---|---|---|---|---|
| rHA Probe | BV421 | - | In House | - | 1:100 |
| FAS | BV510 | Jo2 | BD Biosciences | 563646 | 1:800 |
| CD19 | BV605 | 6D5 | BioLegend | 115540 | 1:800 |
| IgD | BV650 | 11-26c.2a | BioLegend | 405721 | 1:400 |
| IgM | BB700 | II/41 | BD Biosciences | 746099 | 1:200 |
| rHA Probe | AF488 | - | In House | - | 1:100 |
| IgG1 | PE | M1 14D12 | Thermo-Scientific | 12-4015-82 | 1:200 |
| CD27 | PE-Dazzle | LG.3A10 | BioLegend | 124228 | 1:200 |
| CD38 | PE-Cy7 | 90 | BioLegend | 102717 | 1:400 |
| IgG2c | AF647 | polyclonal | Southern Biotech | 1079-31 | 1:200 |
| B220 | AF700 | RA3-6B2 | Biolegend | 103232 | 1:400 |
| TCRb | APC_Fire | H57-597 | Biolegend | 109246 | 1:400 |
| TER-119 | APC_Fire | TER-119 | Biolegend | 116250 | 1:400 |
| Live/Dead | eFluor 780 | - | eBioscience | 65-0865-14 | 1:2000 |

**Table S6. MBC panel, related to Figures 2 and S2.**

| Stain | Fluorochrome | Clone | Vendor | Catalogue # | Dilution |
|---|---|---|---|---|---|
| CD80 (B7-1) | BUV 395 | 2D10.4 | BD | 751728 | 1:200 |
| CD4 | BUV 563 | SK3 | BD | 612912 | 1:200 |
| CD11c | BUV 661 | B-LY6 | BD | 612967 | 1:100 |
| CD141 (BDCA3) | BUV 737 | 1A4 | BD | 741867 | 1:50 |
| L/D | Zombie Aqua | | Biolegend | 423101 | 1:200 |
| CD20 | BV 605 | L27 | BD | 740333 | 1:100 |
| CD11b | BV 650 | ICRF44 | Biolegend | 301336 | 1:100 |
| HLA-DR | BV 711 | G46-6 (aka L243) | BD | 563696 | 1:100 |
| CD303 (BDCA2/CLEC4C) | BV 785 | 201A | Biolegend | 354221 | 1:50 |
| CD16 | PerCP-Cy5.5 | 3G8 | Biolegend | 302027 | 1:50 |
| CD8a | PE-Cy5 | RPA-T8 | Biolegend | 301009 | 1:100 |
| CD1c (BDCA1) | PE-Cy7 | L161 | Biolegend | 331515 | 1:200 |
| CD19 | APC | HIB19 | BD | 555415 | 1:100 |
| CD3 | AF 700 | SP34-2 | BD | 561805 | 1:100 |
| CD14 | APCe780 | 61D3 | Invitrogen | 47-0149-42 | 1:400 |

**Table S7. Human PBMC panel, related to Figures 5 and S5.**

| Stain | Fluorochrome | Clone | Vendor | Catalogue # | Dilution |
|---|---|---|---|---|---|
| CXCR5 | BV421 | L138D7 | BioLegend | 145512 | 1:50 |
| CD19 | BV605 | 6D5 | BioLegend | 115540 | 1:100 |
| IFNg | BV650 | XMG1.2 | BioLegend | 505832 | 1:100 |
| CD4 | PerCP-Cy5.5 | RM4-5 | BioLegend | 100540 | 1:200 |
| CD44 | FITC | IM7 | BioLegend | 103006 | 1:200 |
| CD62L | BUV395 | MEL-14 | BD | 740218 | 1:400 |
| Human IgG | PE | polyclonal | Invitrogen | 12-4998-82 | 1:50 |
| PD-1 | PE-Cy7 | EH12.2H7 | BioLegend | 329918 | 1:200 |
| IL-4 | AF647 | 11B11 | Biolegend | 504110 | 1:100 |
| IL-17A | AF700 | TC11-18H10.1 | Biolegend | 506914 | 1:100 |
| Live/Dead | eFluor 780 | - | eBioscience | 65-0865-14 | 1:2000 |

**Table S8. ICS panel, related to Figures 2 and S2.**

| Stain | Fluorochrome | Clone | Vendor | Catalogue # |
|---|---|---|---|---|
| MHCII | BV421 | M5/114.15.2 | BioLegend | 107632 |
| CD11c | APC | N418 | BioLegend | 117310 |
| CD11b | BV711 | M1/70 | BioLegend | 101242 |
| Anti-Biotin | PE | Bio3-18E7 | Miltenyi Biotec | 130-090-756 |
| anti-CD3e | BV785 | 145-2C11 | BioLegend | 100355 |
| B220 | BV785 | RA3-6B2 | BioLegend | 103245 |
| NK1.1 | BV785 | PK136 | BioLegend | 108749 |

**Table S9. LIPSTIC panel, related to Figures 3 and S3.**
